# A synbiotic medical food improves gut barrier function, reduces immune responses, and inhibits osteoclast activity in models of postmenopausal bone loss aligned with clinical outcomes

**DOI:** 10.1101/2025.08.07.669142

**Authors:** Ryan S. Green, Tyler Roy, Daniela Diaz-Infante Morales, Claire Morrow, Ryan Neilson, Eric M. Schott, Mark R. Charbonneau, Alicia E. Ballok, Katherine J. Motyl, Gerardo V. Toledo

## Abstract

Over half of women above age 50 are affected by osteopenia or osteoporosis, bone-loss conditions influenced by estrogen decline, inflammation, and the intestinal microbiota. Probiotic-based interventions have shown promise in preclinical osteoporosis models. In a recent randomized, double-blind, placebo-controlled clinical trial of postmenopausal women, dietary intervention with SBD111, a synbiotic medical food combining plant-derived probiotics and prebiotic fibers, reduced bone loss in women with osteopenia, elevated body mass index (BMI), and/or elevated body fat.

To investigate potential mechanisms underlying these outcomes, we examined intestinal epithelial, immune, and osteoclast responses to SBD111 in vitro. SBD111 administration improved intestinal barrier integrity, reduced immune cell cytokine secretion, and inhibited osteoclast activity. These effects align with clinically observed reductions in severe gastrointestinal symptoms and bone resorption markers. Together, these findings suggest that SBD111 modulates the gut–bone axis via barrier, immune, and antiresorptive pathways, supporting its role in maintaining skeletal health in postmenopausal women.

**IRB Statements:** The study was conducted according to the guidelines of the Declaration of Helsinki and approved by the Institutional Review Board of MaineHealth under protocol 1689738-1 (approved 12/29/2020) and 958914 (approved 05/31/2005). Human peripheral blood mononuclear cells (PBMCs) were purchased from Charles River Laboratories and collected under their IRB-approved protocol with informed consent for commercial research use.

Informed Consent Statements

Informed consent was obtained from all participants involved in the study.

**Highlights:**

- SBD111 synbiotic medical food improves intestinal barrier function
- SBD111 reduces cytokine release from inflamed immune cells
- This synbiotic medical food inhibits osteoclast activity in vitro
- In vitro SBD111 effects align with clinical reductions in bone loss and CTX-1

**Graphical Abstract.:** **SBD111 synbiotic medical food reduces bone loss via barrier improvement, anti-inflammation, and osteoclast inhibition. (A)** SBD111 improves barrier integrity in vitro. Visualized in the left panel, inflammation reduces intestinal epithelial barrier function, resulting in the influx of inflammatory mediators, thereby activating the immune system. The addition of SBD111, shown in purple (right), improves barrier function, reducing the ability of inflammatory mediators to cross the epithelial barrier. **(B)** SBD111 reduces inflammatory responses of inflamed immune cells in a concentration-dependent manner. Low concentrations of SBD111 induce inflammatory responses. At higher concentrations, fewer inflammatory cytokines are secreted. **(C)** SBD111-derived components/metabolites inhibit the activity of bone-resorbing osteoclasts and reduce their ability to degrade bone. Image created in https://BioRender.com

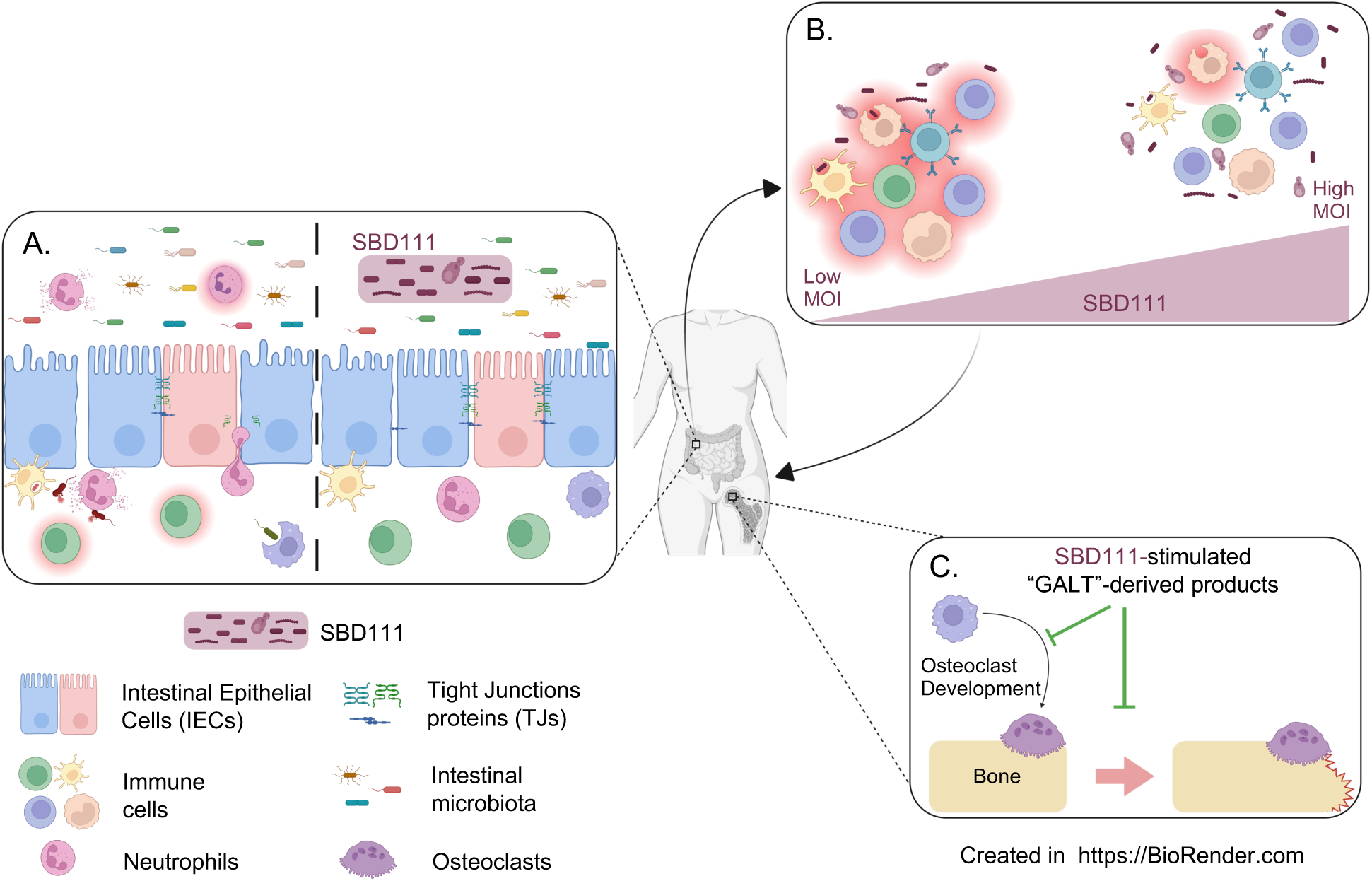

## 1. Introduction

Declining estrogen production during and after menopause is associated with systemic inflammation and rapid bone loss, and it is estimated that 50% of women over the age of 50 have osteoporosis or osteopenia (Greendale et al., 2012; Reid & McClung, 2024; Shieh et al., 2020). These women have a substantially higher risk of hip fracture than those with healthy bone mineral density (BMD), indicating a clear need to manage bone loss during and after menopause (Reid & McClung, 2024). While several pharmaceutical options exist to treat osteoporosis, including hormone therapy and bisphosphonates, adoption of these solutions is limited due to rare but serious side effects, such as increased risk of certain cancers and osteonecrosis of the jaw (Khan et al., 2023; MacLennan et al., 2004; Migliorati et al., 2010). Additionally, options to slow bone loss prior to the development of osteoporosis are limited, as these drugs are not typically prescribed for women with osteopenia.

Menopause-associated bone loss is driven by dysregulation of bone formation and resorption that is characterized by increased osteoclast activity (Møller et al., 2020; Šromová et al., 2023). Classically, osteoclastogenesis is understood to be directed by the receptor activator of nuclear factor-κB ligand (RANKL), macrophage colony stimulating factor (M-CSF), and osteoprotegerin (OPG) (Boyle et al., 2003; Takegahara et al., 2024). In addition to these factors, inflammatory cytokines, including interleukin-6 (IL-6), IL-23, and interferon gamma (IFN-γ), have been shown to modulate osteoclast activity and osteoclastogenesis (Cai et al., 2024; S. Li et al., 2024; Umur et al., 2024).

Thus, it is now recognized that systemic inflammation plays a key role in peri- and postmenopausal bone loss (Ferbebouh et al., 2021; Zhang et al., 2022).

Substantial evidence indicates that systemic inflammation and skeletal health are regulated, in part, by the gut microbiota, a diverse community of microbes that inhabits the human gastrointestinal (GI) tract (J. Y. Li et al., 2016; Mazziotta et al., 2023; Zaiss et al., 2019). For example, gut microbes can modulate integrity of the GI epithelial barrier that serves as a primary defense against pathogens and inflammatory insults (Di Vincenzo et al., 2023a; Thoo et al., 2019). This barrier is impaired by menopause, aging, and obesity, allowing microbe-derived compounds to enter the lamina propria, induce the production of inflammatory cytokines and chemokines, and increase systemic inflammation (Di Vincenzo et al., 2023a; Shieh et al., 2020; Teixeira et al., 2012; Thevaranjan et al., 2017; Thoo et al., 2019). Certain microbial metabolites, including the short chain fatty acids butyrate and acetate, can directly inhibit these inflammatory processes and improve epithelial barrier integrity (Furusawa et al., 2013; Hosmer et al., 2024). Microbial metabolites have also been implicated as regulators of musculoskeletal health, inhibiting bone resorption by osteoclasts directly and indirectly through immunological mechanisms (Rahman et al., 2003; Tyagi et al., 2018; Zaiss et al., 2019). The structure and function of the human intestinal microbiota are substantially influenced by diet. These observations suggest that dietary interventions targeting the gut microbiota could be developed to regulate GI barrier function, systemic inflammation, and bone loss in peri- and postmenopausal women (Moles & Otaegui, 2020).

To address the unmet need of menopause-associated bone loss, we developed SBD111, a synbiotic formulation designated as a medical food for the dietary management of postmenopausal bone loss. SBD111 is comprised of four probiotic strains derived from fruits and vegetables: *Levilactobacillus brevis*, *Lactiplantibacillus plantarum*, *Leuconostoc mesenteroides*, and *Pichia kudriavzevii,* as well as prebiotic fibers, oligofructose and blueberry powder, which can serve as growth substrates for these organisms (Easson et al., 2022; Lawenius et al., 2022). This synbiotic was formulated to synergistically produce acetate and to deliver a much higher concentration of viable microbes than traditional probiotic foods or supplements while including prebiotic fibers to further enhance microbial viability and function (Pandey et al., 2015).

The present study was motivated by findings from a recent prospective, multicenter, double-blind, randomized, placebo-controlled efficacy trial of SBD111 in 286 healthy women within six years of menopause (Clinical Trial ID Number: NCT05009875) (Schott et al., 2025). In this trial, SBD111 reduced bone loss in two prespecified populations: women with osteopenia and those with a body mass index (BMI) ≥30. A similar reduction was also observed in a post-hoc analysis of women with ≥40% body fat (Schott et al., 2025). Furthermore, in women with elevated BMI, SBD111 administration was associated with decreased serum concentrations of collagen cross-linked telopeptide (CTX-1; a marker of bone degradation), suggesting inhibition of osteoclast function (Schott et al., 2025). These findings are notable, as osteopenia, elevated BMI, and elevated body fat are associated with increased systemic inflammation (Ferbebouh et al., 2021; Festa et al., 2001; Zhang et al., 2022). These observations also align with preclinical data wherein administration of a preliminary SBD111 formulation in an ovariectomized mouse model of menopausal-bone loss significantly reduced trabecular bone loss and expression of the inflammatory cytokines *Tnf* (encoding tumor necrosis factor alpha [TNF-α]) and *Il6* (encoding IL-6) within vertebral bone, further implicating that immunological regulation underlies SBD111s function (Lawenius et al., 2022).

Given the known links between inflammation, adiposity, and osteoclast activity, we hypothesized that SBD111 modulates the gut–immune–bone axis through epithelial, immunological, and resorptive pathways.

Here, we report that SBD111 administration improves intestinal epithelial barrier integrity, elicits concentration-dependent anti-inflammatory responses, and inhibits osteoclast activity in vitro. Each of these putative mechanisms has the potential to synergistically reduce bone loss in postmenopausal women with osteopenia or elevated BMI following administration of SBD111.

## 2. Methods

### 2.1 Microbial Strains and Preparation

SBD111, a synbiotic composed of lyophilized *Levilactobacillus brevis*, *Lactiplantibacillus plantarum*, *Leuconostoc mesenteroides*, *Pichia kudriavzevii*, and prebiotic fibers, has been previously described (Easson et al., 2022; Sahni et al., 2023). Previous formulations of SBD111 (SBD111-A) included *P. fluorescens,* which was removed from the final formulation of SBD111, due to low acetate production (Lawenius et al., 2022).

SBD111 material was resuspended at a concentration of 1.48 x 10^9^ total colony forming units (CFU)/mL (CFU/mL by strain: 7.81 x 10^7^ CFU/mL *P. kudriavzevii*; 4.69 x 10^8^ CFU/mL of each: *L. brevis, L. mesenteroides,* and *L. plantarum*) and ∼9.38 mg/mL (capsule to capsule variation within a range of 8.13 – 11.25 mg/mL) of each prebiotic component and in 1 X phosphate buffered saline (PBS; Catalog (CAT)# BP3991, Thermo Fisher; Waltham, MA) with regular mixing for 25 min at room temperature.

*Escherichia coli* (Strain: 1100101; CAT# BAA-2471, American Type Culture Collection (ATCC), Manassas, VA) was grown overnight at 37°C in Tryptic Soy Broth (TSB; CAT# 1.00800.0500, Merk KGAG; Darmstadt, Germany). *E. coli* was washed with 1 X PBS and resuspended in Minimum Essential Medium (MEM, phenol red-free; CAT# 51200038, Gibco^TM^, Thermo Fisher; Waltham, MA), supplemented with 10% fetal bovine serum (FBS; CAT# 16140071 Gibco^TM^, Thermo Fisher; Waltham, MA) and 1 X Glutamax^TM^ (CAT# 35050061, Gibco^TM^, Thermo Fisher; Waltham, MA), to a concentration of 2.5 x 10^7^ CFU/mL.

### 2.2 Cells and Culture Conditions

Caco-2 human adenocarcinoma cells (CAT# 86010202, European Collection of Authenticated Cell Cultures, MiliporeSigma; Merck KGAG, Darmstadt, Germany) and RAW264.7 murine macrophages (CAT# TIB-71, ATCC; Manassas, VA) were cultured at 37°C and 5% CO_2_ in Dulbecco’s Modified Eagle Medium (DMEM; CAT# 10566016, Gibco^TM^, Thermo Fisher; Waltham, MA) supplemented with 10% FBS, 1 X antibiotic-antimycotic (anti-anti; CAT# 15240062, Gibco^TM^, Thermo Fisher; Waltham, MA), and 1 X Glutamax^TM^. HT29-Lucia^TM^ AHR cells (CAT# ht2l-ahr, Invivogen; San Diego, CA) human adenocarcinoma cells were grown at 37°C and 5% CO_2_ in DMEM supplemented with 10% FBS, 1 X Glutamax^TM^, 1 X anti-anti, and 100 mg/mL Zeocin® (CAT# ant-zn-05, Invivogen; San Diego, CA). Cell lines were used from passage 3 to 15 for described assays. Human peripheral blood mononuclear cells (PBMCs) were purchased from Charles River Cell Solutions (CAT# PB009C-2, Northridge, CA). Purchased PBMCs were thawed and used immediately upon receipt. PBMCs were cultured at 37°C and 5% CO_2_ in Roswell Park Memorial Institute Medium (RPMI-1640) without phenol red (CAT# 11835030, Gibco^TM^, Thermo Fisher; Waltham, MA) supplemented with 10% FBS, 1 X Glutamax^TM^, and 12.5 mM HEPES Buffer (CAT# 15630130, Gibco^TM^, Thermo Fisher; Waltham, MA). PBMCs were maintained in culture for no more than 32 h.

### 2.3 Barrier integrity assay

Caco-2 and HT29 cells were cultured separately and combined at a 70:30 ratio of Caco-2 and HT29 cells, respectively (Ferraretto et al., 2018). A total of 2.5 x 10^4^ mixed Caco-2 and HT29 cells were seeded per insert on polycarbonate membrane cell culture inserts (6.5 mm diameter, 0.4 µm pore size; 3413, CAT# CLS3396-2EA Costar^TM^; Corning, NY), as depicted in **Fig.1A**. Each insert contained 200 µL of DMEM in the apical chamber, while the basolateral chamber received 1 mL of the same medium. The plates were maintained at 37°C with 5% CO_2_. The day before the assay, the inserts were washed with 1 X PBS, and the culture medium was replaced with an antibiotic-free medium: phenol red-free MEM, supplemented with 10% FBS and 1 X Glutamax^TM^, with 180 µL added to the apical chamber and 1 mL to the basolateral chamber.

**Figure 1.**
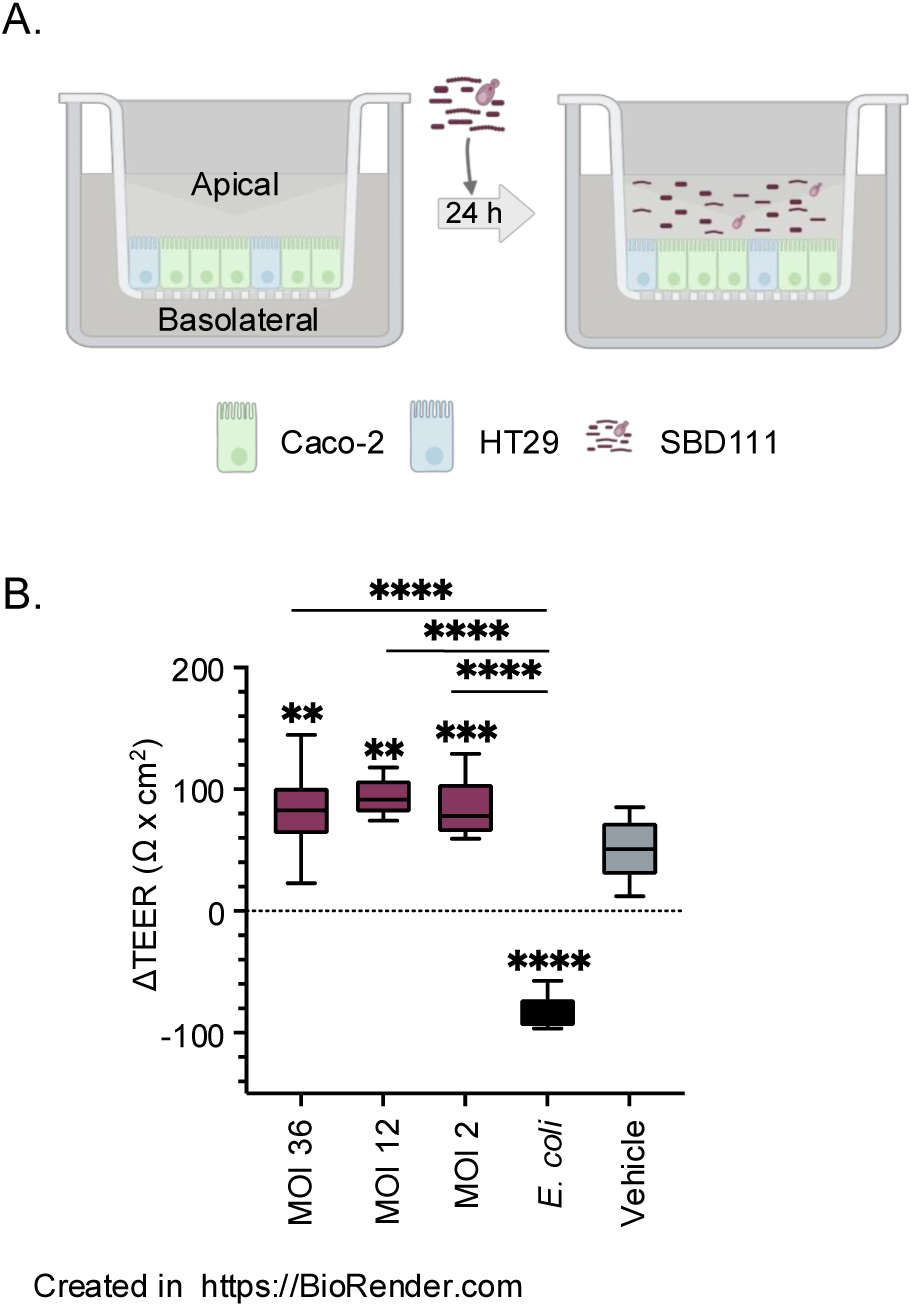
SBD111 increases barrier function (TEER) of intestinal epithelial monolayers. **(A)** Mature, polarized intestinal epithelial cell (Caco-2 and HT29) monolayers were exposed to a media control (Vehicle), disruption control (*E. coli*), or SBD111 material at a multiplicity of interaction (MOIs; microbes per human cell) of 36, 12, or 2 for 24 h. Graphic created in https://BioRender.com. **(B)** Trans-epithelial electrical resistance (TEER) was measured before and after exposure to SBD111 (MOI 36, 12, and 2) for each monolayer, and the mean change in TEER (DTEER) over time was compared across the different conditions. Data presented is a single experiment that is representative of eight total experiments. The values plotted represent the means of 12 replicates, with boxes representing interquartile range and error bars indicating the minimum and maximum values. Significance between conditions was determined by one-way ANOVA with Tukey’s HSD. Asterisks without comparison bars indicate significance relative to the vehicle control. (** = *p*<0.005, *** = *p*<0.001, **** = *p*<0.0001)

On days 18 to 21, the trans-epithelial electrical resistance (TEER) of Caco-2-HT29 cell monolayers was measured using a voltohmmeter (Millicell® ERS-2; CAT# MERS00002, EMD Millipore Corporation, Burlington, MA). A pair of electrodes (MERSSTX01, EMD Millipore Corporation, Burlington, MA) was inserted into each well to measure the resistance. TEER measurements were taken before the experiment (T0) to establish a baseline and 24 h after microbial treatment (T1) to assess the change in TEER (ΔTEER). The values were corrected for background resistance and expressed as ν x cm². The average TEER values ranged from 150 to 250 ν x cm².

Apical Caco-2:HT29 monolayers were co-incubated with 20 µL of either resuspended SBD111 material or a disruption control. Resuspended SBD111 material was diluted in MEM antibiotic-free medium to a multiplicity of interaction (MOI, microbes per human cell) of 36, 12, 2, or 0. Disruption controls were *E. coli* diluted in anti-biotic free MEM to an MOI of 20 (2.5 x 10^7^ CFU/mL). The plates were incubated for 24 h at 37°C with 5% CO_2_. The apical and basolateral supernatants were collected separately for cell viability and cytokine analysis.

### 2.4 SBD111 treatments of PBMCs

Cryopreserved human PBMCs representing seven donors (four peri- and postmenopausal females, ages 51-59 years with an average age of 55 years, and three males, age 22-50 years with an average age of 37 years) were purchased from Charles River Cell Solutions (CAT# PB009C-2, Northridge, CA). In 1.5 mL, one million cells were treated with media alone (vehicle control), 100 ng/mL of lipopolysaccharide (LPS, CAT# tlr-eblps, Invivogen; San Diego, CA) in RPMI, or resuspended SBD111 capsule contents diluted in RPMI to a MOI of 10 or 1 (C. Kleiveland & Kleiveland, 2015). Cells were incubated for 24 h at 37°C with 5% CO_2_; after which supernatants were harvested for viability and cytokine response analysis. Cell viability was determined via lactate dehydrogenase activity as per manufacturer’s instructions (CAT# G1780, Promega; Fitchburg, WI). Cytokines were analyzed via ELISA. ELISAs were performed as per manufacturer’s instructions (Thermo Fisher, Waltham, MA: TNF-α (CAT# 88-7346-88), IL-23p19 (CAT# 88-7237-88); BioLegend, San Diego, CA: IL-12p70 (CAT# 431701), IL-6 (CAT# 430501), IL-1β (CAT# 437016), IL-8/C-X-C motif ligand 8 (CXCL8) (CAT# 431501); and R&D systems, Minneapolis, MN: IFN-γ (CAT# DY285B), CXCL1 (CAT# DY275), OPG (CAT# DY805), IL-10 (CAT# DY217B)).

### 2.5 PBMC LPS challenge and SBD111 treatments

Frozen PBMCs from the healthy donors described above were incubated at 37°C with 5% CO_2_ for 30 min with either 100 ng/mL of LPS (CAT# tlr-eblps, Invivogen; San Diego, CA) in RPMI, or with media alone (unchallenged control) (C. Kleiveland & Kleiveland, 2015; Ngkelo et al., 2012). After incubations, cells were washed once with 1 X PBS and resuspended in RPMI. In 1.5 mL, one million LPS-challenged or unchallenged cells were treated with an RPMI control (Vehicle) or resuspended SBD111 material diluted in RPMI to a MOI of 10 or 1. Cells were incubated for 24 h at 37°C with 5% CO_2_. Resulting supernatants were harvested and examined for viability and cytokine responses. Cell viability was determined via lactate dehydrogenase activity and cytokines were analyzed via ELISA as described above.

### 2.6 Production of GALT model supernatants

1 x 10^5^ HT29 and Caco-2 cells were seeded onto polycarbonate membrane cell culture inserts (12 mm diameter, 0.4 µm pore size; CAT#3401, Costar; Corning, NY) at a 70:30 ratio of Caco-2 to HT29 cells in 0.5 mL of DMEM (Ferraretto et al., 2018). 1.5 mL of DMEM was added to each well. After 1 week of culture media was replaced with αMEM without phenol red (CAT#41061029, Gibco^TM^, Thermo Fisher; Waltham, MA) supplemented with 10% charcoal-stripped FBS (CAT# F6765, MiliporeSigma; Merck KGAG, Darmstadt, Germany) and 1 X Glutamax^TM^ for an additional 11-14 days; during which media continued to be replaced every 48 h. Cryopreserved PBMCs from a peri-, postmenopausal female donor (age 58 years) were thawed and resuspended in αMEM supplemented with 10% charcoal-stripped FBS and 1 X Glutamax^TM^. One million PBMCs were added to the basolateral chamber of each epithelial cell monolayer containing well to produce a gut-associated lymphoid tissue (GALT) model as described previously (C. R. Kleiveland, 2015; Korsten et al., 2023). The apical side of each GALT model was incubated with an αMEM control (vehicle), SBD111 material, diluted to an MOI of 10 or 2 (relative to epithelial barrier), or LPS, diluted to 500 ng/mL in αMEM supplemented with 10% charcoal-stripped FBS and 1 X Glutamax^TM^. Cells were incubated for 24 h at 37°C with 5% CO_2_; after which basolateral supernatants were harvested for viability, osteoclastogenesis modeling, and cytokine response analysis.

Supernatants harvested for osteoclastogenesis modeling were filter sterilized with 0.2 µm cellulose-acetate syringe filters. Cells were harvested for RANKL and OPG gene expression analysis via qRT-PCR.

### 2.7 RAW264.7 cell osteoclast model

2.5 x 10^4^ RAW264.7 macrophages in 140 µL of DMEM were incubated for 7 days at 37°C with 5% CO_2_ in DMEM containing 50 ng/mL of RANKL (Quach et al., 2019). Cells were treated with 60 µL of undiluted basolateral supernatants from the GALT models, described above. Alternatively, RAW264.7 macrophages were treated with SBD111-conditioned αMEM, which was produced by culturing resuspended SBD111 material at a concentration of 6.67 x 10^5^ or 1.33 x 10^5^ CFU/mL (MOI 10 and 2 equivalent concentrations) in αMEM (supplemented with 10% charcoal-stripped FBS and 1 X Glutamax^TM^) for 24 h at 37°C and 5% CO_2_. The resulting conditioned supernatant was filter sterilized with 0.2 µm cellulose-acetate syringe filters prior to incubation with RAW264.7 cells. After one week, supernatants were harvested to assay lactate dehydrogenase activity as per manufacturer’s instructions to determine viability (CAT# G1780, Promega; Fitchburg, WI). Cells were analyzed for tartrate-resistant acid phosphatase (TRAP) activity to quantify osteoclast differentiation as per manufacturer’s instructions (CAT# MK301, Takara; San Jose, CA).

### 2.8 Human osteoclast model

Human PBMCs were isolated from whole blood samples collected from four female patients ages (65-87) mean age of 74 into EDTA tubes. Blood samples were transferred into a 50 mL centrifuge tube (CAT# 21008-690, Corning; Corning, NY) and an equal volume of sterile 1 X PBS Buffer with 2% FBS (CAT# 07905, Stemcell^TM^ Technologies; Cambridge, MA) was added. PBMCs were separated using Sepmate^TM^ tubes (CAT# 85460, Stemcell^TM^ Technologies; Cambridge, MA) with 15 mL Ficoll-Paque^TM^ Plus (CAT# 95021-205, Cytiva; Marlborough, MA) and centrifuged for 20 min at 1200 *x g*.

Isolated PBMCs were decanted into a new 50 mL centrifuge tube washed with 40 mL of 1 X PBS with Buffer 2% FBS and centrifuged at 300 *x g* for 10 min. Following centrifugation, the supernatant was discarded, and wash steps were repeated for a second time. Finally, PBMC pellet was triturated and suspended in plating media on ice and counted manually with a hemocytometer.

#### 2.8.1 Human osteoclast culture

PBMCs isolated from human blood samples were seeded on bone chips (CAT# DT-1BON1000-96, Immunodiagnostic Systems, East Boldon, UK; Boneslices.com) at a density of 1.25 x 10^6^ cells/cm^2^ which is equivalent to 4 x 10^5^ cells/well on a 96 well plate. For the culture, PBMCs and bone chips are suspended in a 2:1 ratio of osteoclastogenic media and GALT supernatant, as produced above. The osteoclastogenic media is comprised of αMEM without phenol red (CAT#41061029, Gibco^TM^, Thermo Fisher; Waltham, MA) supplemented with 10% charcoal-stripped FBS (CAT# F6765-500ML, MiliporeSigma, Merck KGAG, Darmstadt, Germany), 1 X Pen-Strep (CAT# 15140122, Thermo Fisher; Waltham, MA), RANKL (CAT# 390-TN-010/CF, R&D systems, Minneapolis, MN), M-CSF (CAT# 216-MC-010, R&D systems, Minneapolis, MN). The media mixture resulted final concentrations of 50 ng/mL RANKL and 25 ng/mL M-CSF. Once PBMCs were seeded they were cultured for 14 days at 37°C with media changes every 48-72 h. Finally, cells were fixed using 2.5% glutaraldehyde (CAT# 16220, Electron Microscopy Sciences; Hatfield, PA) on day 14 after conditioned media was collected.

#### 2.8.2 Quantifying osteoclast differentiation and activity

Media was collected from day 12 of the osteoclast culture to measure CTX-1 released into media as a measure of osteoclast resorptive activity. Levels of CTX-1 in each well of treated media were analyzed using CrossLaps® for Culture (CTX-I) ELISA (CAT# AC-07F1, Immunodiagnostic Systems, East Boldon, UK).

On day 14 cells were fixed with 2.5% glutaraldehyde, and the bone chips were stained for TRAP positive cells using a TRAP kit (CAT # 387A-1KT, MiliporeSigma, Merck KGAG, Darmstadt, Germany) for 1 h and 45 min. Stained bone chips were then imaged using the Keyence BZ-X800 microscope at 10x magnification. TRAP positive osteoclasts were quantified, and total area was analyzed using the NOISe machine learning algorithm (Kumar et al., 2024).

### 2.9 Statistical analyses

Data were analyzed using the Prism graphing and analysis software (GraphPad Software, Boston, MA). Significance was defined as *p* < 0.05 as determined using one-way ANOVA with Tukey’s HSD. When multiple donors were examined one-way ANOVA with Tukey’s HSD was performed within each donor.

## 3. Results

### 3.1 SBD111 administration improves intestinal barrier integrity

SBD111 administration was associated with reduced severe gastrointestinal symptoms in post-menopausal women, indicating direct effects on cells of the GI tract. Additionally, as SBD111 is administered orally in an enterically-coated capsule format and initially interacts with the human body at the intestinal epithelium, we prioritized investigating the effects of SBD111 on intestinal epithelial cells. To examine whether SBD111 affects barrier integrity, intestinal epithelial cell monolayers were established on semi-permeable membranes, as illustrated in **Fig. 1A**. These monolayers were exposed to a media control (Vehicle), or to SBD111 material, including prebiotic fibers, at a total MOI of 36, 12, or 2. Entire capsule contents were utilized as prebiotics are potentially immunologically active and can alter SBD111 viability, growth, and production of beneficial metabolites. Additionally, as SBD111 is orally delivered in an enterically coated capsule, intestinal cells interact with SBD111 at varying concentrations depending on their location in GI tract. As approximately 10 bacteria can directly bind to an individual intestinal epithelial cell, the SBD111 MOI range utilized in this study represents concentrations that human intestinal epithelial cells are expected to encounter after capsule release (Dimitrov et al., 2014). *E. coli* was used as a barrier disruption control, as it has been demonstrated to damage barrier integrity in vitro (Yuan et al., 2020). The effect of SBD111 on intestinal barrier function was assessed by changes in TEER across the monolayer at 0 and 24 h incubation. As shown in **Fig. 1B**, all tested MOIs — 36, 12, and 2 — resulted in a significant increase in TEER compared to the vehicle control, though there were no significant differences between these groups. *E. coli* significantly reduced TEER as expected.

Intestinal epithelial cell immune responses toward SBD111 were examined through the secretion of IL-8 and CXCL1, chemokines associated with intestinal inflammation and neutrophil recruitment (**Supplementary** Fig. 1) (Capucetti et al., 2020). SBD111 administration did not significantly increase epithelial chemokine secretion compared to the vehicle control. By contrast, secretion of both chemokines was significantly increased by *E. coli* application.

### 3.2 SBD111 elicits concentration-dependent anti-inflammatory responses from human immune cells

To explore how SBD111 may modulate systemic inflammation to benefit populations with osteopenia or high BMI, we treated human peripheral blood mononuclear cells (PBMCs) with increasing concentrations of SBD111. This approach was motivated by the known association between elevated BMI, bone loss, and inflammation, and by preclinical signals suggesting inflammatory modulation in treated participants (Ferbebouh et al., 2021; Festa et al., 2001; Lawenius et al., 2022; Schott et al., 2025; Zhang et al., 2022). Cryopreserved human PBMCs isolated from four peri- or postmenopausal female donors as well as three male donors were treated with a media control (vehicle), a stimulatory control (LPS), or SBD111 material at a total MOI of 10 or 1. It has been established that some immune cells directly monitor the intestinal lumen and that specialized cells translocate bacteria into immunological compartments (Da Silva et al., 2017; Farache et al., 2013). For this reason, an MOI of 1 was chosen to represent the low abundance stimulation experienced by intestinal immune cells in a healthy gut. Additionally, a higher MOI of 10 was included to reflect the increased bacterial translocation into tissues that accompanies the menopause transition and high BMI/body fat, a population that benefits from SBD111 administration (Shieh et al., 2020; Teixeira et al., 2012; Thevaranjan et al., 2017). After 24 h, cytokine responses were analyzed. As expected, we found that SBD111 stimulated inflammatory cytokine secretion in inflammation-naïve PBMCs **(Fig. 2A, C, E, G, and I)**. As PBMCs protect the body from microbial challenge, this response is anticipated for probiotic microbes (Hua et al., 2010; Ren et al., 2019). However, an SBD111 concentration-dependent reduction in cytokine secretion was observed for many inflammatory cytokines. Across the four female donors, SBD111 administration at an MOI of 10 resulted in significantly reduced secretion of IL-23, IL-8, CXCL1, and IFN-γ relative to an MOI of 1 **(Fig 2A, E, G, and I)**. These cytokines are important inflammatory signals that function in inflammatory T cell polarization/maintenance (IL-23 and IFN-γ) and neutrophil chemoattraction (IL-8 and CXCL1) (Capucetti et al., 2020; Cui et al., 2025; Muranski & Restifo, 2013). For CXCL1 and IFN-γ, SBD111 administration at an MOI of 10 did not result in a significant increase compared to the media control. The three male donor-derived PBMCs responded similarly to the female donors except for IL-23 and IL-8 secretion, which exhibited the same trend as the female donors but was statistically insignificant **(Fig. 2A and E)**. It is notable that donor variation was seen across cytokine responses, however each donor trended similarly, as indicated by donor specific coloration in **Fig. 2** and shown as absolute cytokine secretion by donor in **Supplementary** Fig. 2 and 3. Other examined cytokines included TNF-a and IL-1b, which were secreted in response to SBD111(**Supplementary** Fig. 4A, B, C, and **D).** IL-12, a Th1 cytokine mirrored the trends seen for IFN-γ, but was highly donor dependent **(Supplementary** Fig. 2F and 3F). The anti-inflammatory factor, IL-10, was secreted variably between donors and was significantly reduced by SBD111 in the inflammatory challenge model (**Supplementary** Fig. 4E and F).

**Figure 2.**
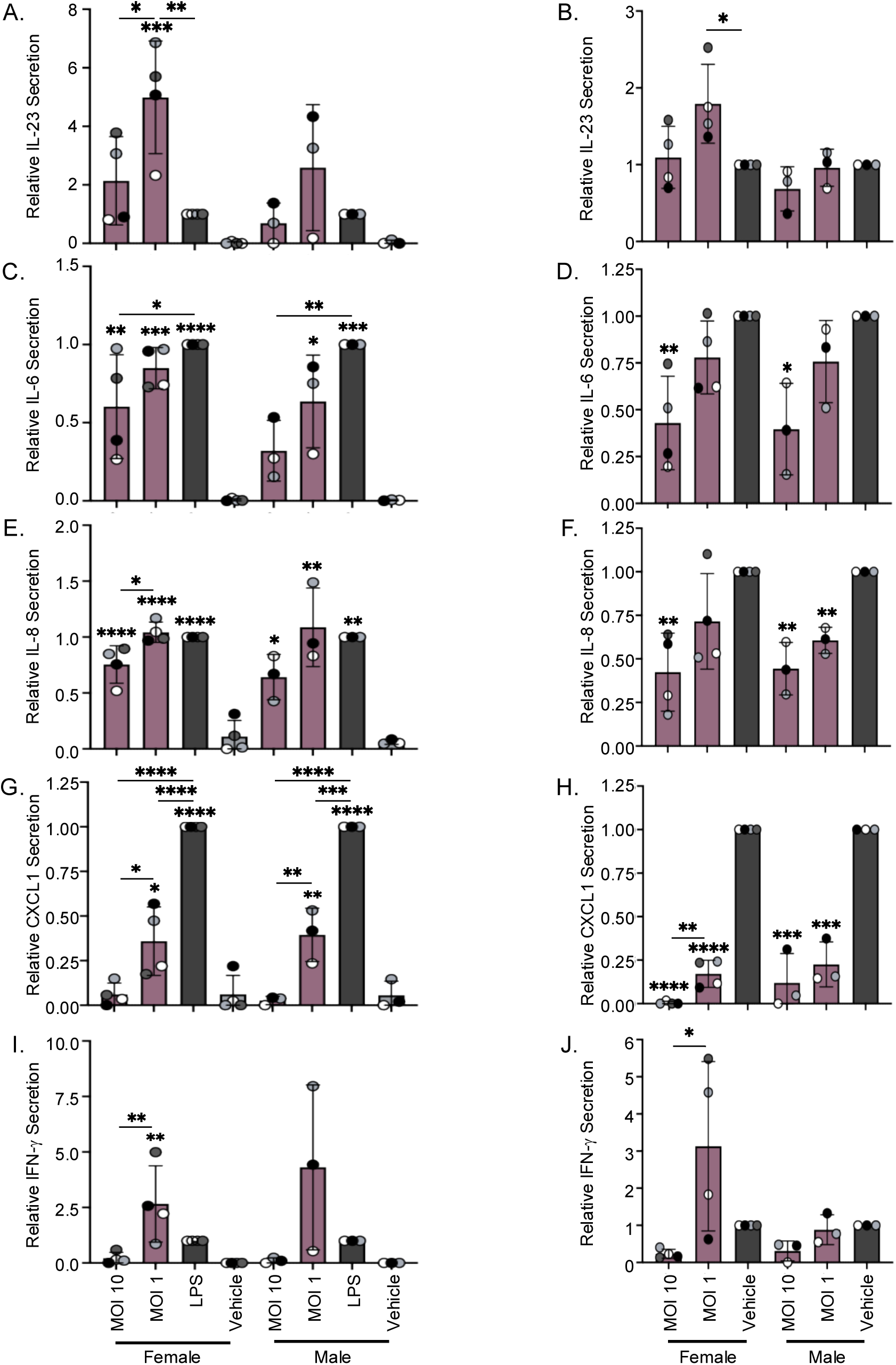
SBD111 induces concentration-dependent reductions in inflammatory responses by human PBMCs both at baseline and after inflammatory challenge. (A, C, E, G, and. **I)** Human peripheral blood mononuclear cells (PBMCs) from four peri- and postmenopausal females and three males were incubated with a vehicle control (media), lipopolysaccharide (LPS; a stimulatory control), or SBD111 material at an MOI of 10 or 1 for 24 h. After the incubation, immune responses elicited against SBD111 were determined. **(B, D, F, H, and J)** Human PBMCs from the same four peri- and postmenopausal females and three males were treated with 100 ng/mL of LPS for 30 min to induce inflammatory responses then washed with 1 X PBS and resuspended in cell culture media. These inflamed cells were then incubated with a vehicle control (media) or SBD111 material at an MOI of 10 or 1 for 24 h. Cytokine responses were quantified via ELISA, **(A and B)** IL-23, **(C and D)** IL-6, **(E and F)** IL-8, **(G and H)** CXCL1, and **(I and J)** IFNγ, and normalized to LPS (inflammation naïve conditions) or LPS-challenged, vehicle-treated controls (inflammatory challenge) to compare across donors and experiments. Columns indicate the mean + SD for all donors, differentiated by sex. Points indicate the mean response of an individual donor and are color coded by donor. Significance between conditions within each sex was determined by one-way ANOVA with Tukey’s HSD. Asterisks without comparison bars indicate significance relative to the vehicle control. (* = *p*<0.05, ** = *p*<0.005, *** = *p*<0.001, **** = *p*<0.0001).

To model the effects of SBD111 administration under conditions more representative of the systemic inflammation observed in osteopenic and BMI 30+ women post-menopause, an LPS challenge model was used (C. Kleiveland & Kleiveland, 2015; Ngkelo et al., 2012). PBMCs isolated from the same four female and three male donors were briefly treated with LPS, after which the LPS was removed and PBMCs were exposed to a media control (vehicle) or SBD111 material diluted to a total MOI of 10 or 1. Brief LPS challenge alone did not robustly induce Th1 cytokine secretion (IFN-γ and IL-12), consistent with prior reports **(Supplemental Fig. 3E and F**) (Janský et al., 2003). In contrast to the inflammation-naïve responses, SBD111 significantly reduced IL-6, IL-8, and CXCL1 secretion in a concentration-dependent manner relative to the LPS-challenged vehicle control in female and male PBMCs **(Fig. 2D, E, and H).** When IL-23 and IFN-γ were examined, both trended similarly to the unchallenged responses for females, however, IL-23 secretion was not significantly reduced by SBD111 at MOI 10 **(Fig 2B and J)**. Additionally, during stimulatory conditions male PBMCs exhibited different trends from female donors for IL-23 and IFN-γ, this is not unexpected as important sources of these cytokines: dendritic cells, monocytes, and T cells, are differ between sexes (Pal et al., 2024; Scotland et al., 2011; White et al., 2022)

### 3.3 SBD111 reduces osteoclast activity

Orally administered Microbes can exert potent immune effects locally in the GI tract that translate into systemic changes, including effects on osteoclastogenesis (Di Vincenzo et al., 2023a; Zaiss et al., 2019). Given the significant reductions is bone loss and serum CTX observed in participants with BMI ≥30 in the clinical trial, we sought to investigate whether SBD111 alters osteoclastogenic signaling via its interaction with GALT in conditions mimicking an intact gut epithelial barrier (C. R. Kleiveland, 2015; Korsten et al., 2023). This model contains many of the relevant cell types found within GALT producing a more physiologically relevant model than individual cell models and has been used to dissect mechanism of action in vitro.

Briefly, intestinal epithelial monolayers were established on cell culture inserts as illustrated in **Fig. 1A**. PBMCs were added to the basolateral chamber to represent the immune components of GALT, while SBD111 was applied to the apical compartment, representing the intestinal lumen, as represented in **Fig. 3A**. After 24 h, the basolateral supernatant (PBMC and intestinal epithelial cell-conditioned supernatant) was harvested to examine the osteoclastogenic potential of secreted factors, and cellular RNA was isolated to quantify gene expression. The expression of pro- and anti-osteoclastogenic factors (*RANKL* and *OPG*, respectively) by the GALT model intestinal epithelial cells and PBMCs was quantified via qRT-PCR. Across four select donors, no significant differences were seen in OPG and RANKL expression across donor PBMCs as a function of SBD111 administration (**Supplementary** Fig. 5).

**Figure 3.**
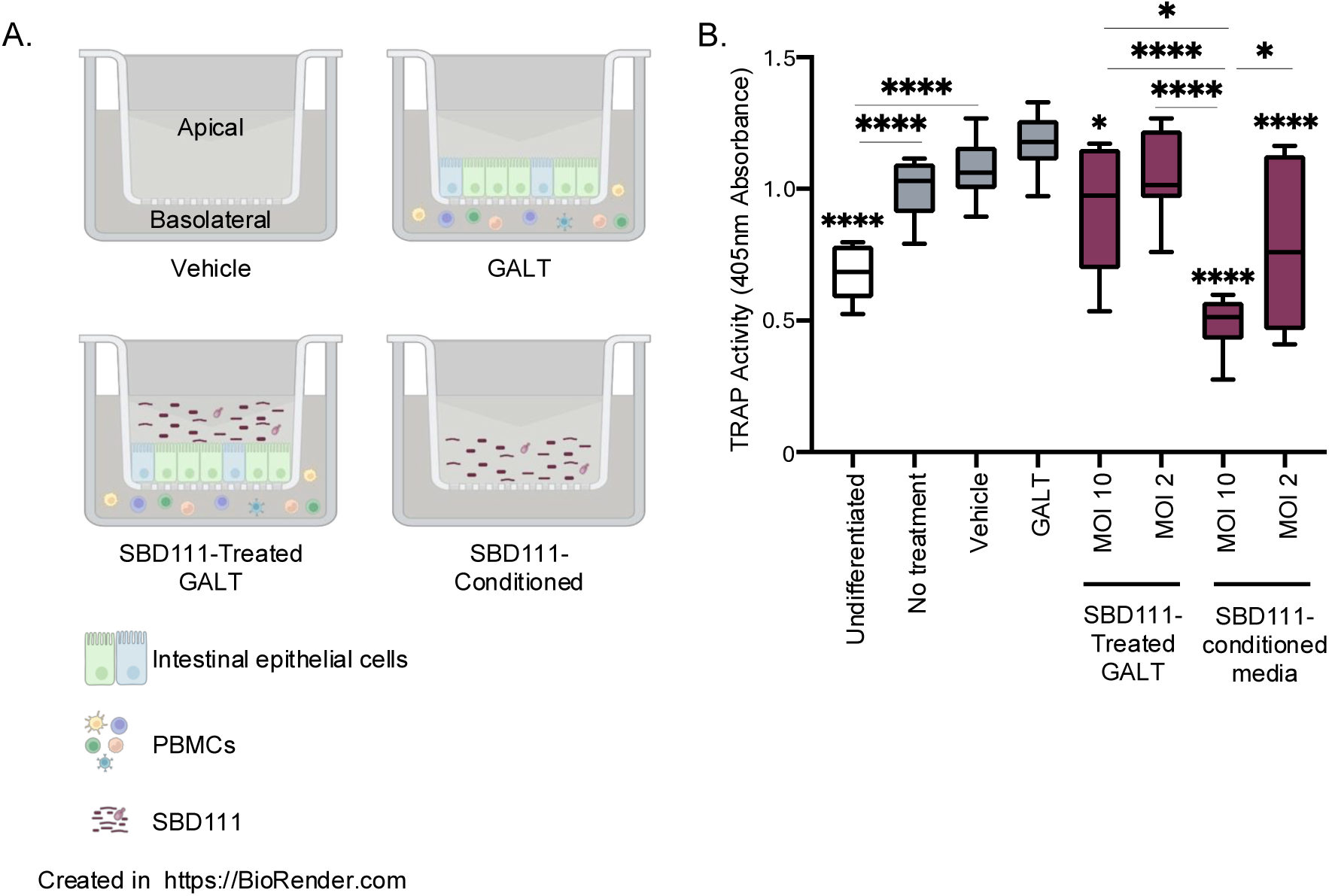
SBD111 reduces osteoclastogenesis in murine RAW264.7 models. Gut-associated lymphoid tissue (GALT) models were established containing polarized intestinal epithelial cell (Caco-2 and HT29) monolayers on cell culture inserts. Human PBMCs were added to the basolateral compartment. A media control (vehicle) or SBD111 material at MOIs of 10 or 2 were added to the apical compartment of the GALT model for 24 h. For SBD111-conditioned media without human cells, concentrations equivalent to an MOI of 10 or 2 were used. **(A)** A graphic representation of the treatments that were used in this experiment. Basolateral supernatants were harvested from each of these conditions to examine their effects on osteoclastogenesis. Graphic created in https://BioRender.com. **(B)** RAW264.7 cells were treated with media (Undifferentiated) or 50 ng/mL of receptor activator of nuclear factor-κB (NF-kB) ligand (RANKL). RANKL-treated cells were further treated with RAW264.7 media (No treatment), GALT model media (Vehicle), media control-treated GALT-conditioned media (GALT), SBD111-treated GALT-conditioned media (MOI 2 or 10), or SBD111-conditioned media without human GALT (MOI 10 or 2). RAW264.7 cells were cultured for seven days. After seven days, osteoclastogenesis was determined by measuring tartrate-resistant acid phosphatase (TRAP) activity. Results are presented as the mean of ³8 data points, with boxes representing interquartile range and error bars indicating the minimum and maximum values. Significance between conditions was determined by one-way ANOVA with Tukey’s HSD. Asterisks without comparison bars indicate significance relative to the GALT control. (* = *p*<0.05, **** = *p*<0.0001).

To characterize the contribution of SBD111-GALT interactions and SBD111 itself to osteoclastogenesis, media controls (Vehicle), GALT-conditioned supernatants (GALT), SBD111-treated GALT-conditioned supernatants (MOI –10 or 2) from the previous section, or SBD111-conditioned supernatants (human cell free, equivalent MOI – 10 or 2; 6.67 x 10^5^ or 1.33 x 10^5^ CFU/mL, respectively) were harvested and applied along with RANKL to RAW264.7 cells, a murine cell line commonly used to study osteoclastogenesis(Quach et al., 2019). Graphic representations of each GALT condition are indicated in **Fig. 3A**. We observed that SBD111-treated GALT-conditioned media reduced TRAP activity in a SBD111 concentration-dependent manner **(Fig. 4B).** SBD111-conditioned media alone, without GALT-derived factors, induced a similar concentration-dependent reduction in osteoclast development. These data indicate that SBD111 secretes components/factors that reduce osteoclastogenesis.

**Figure 4.**
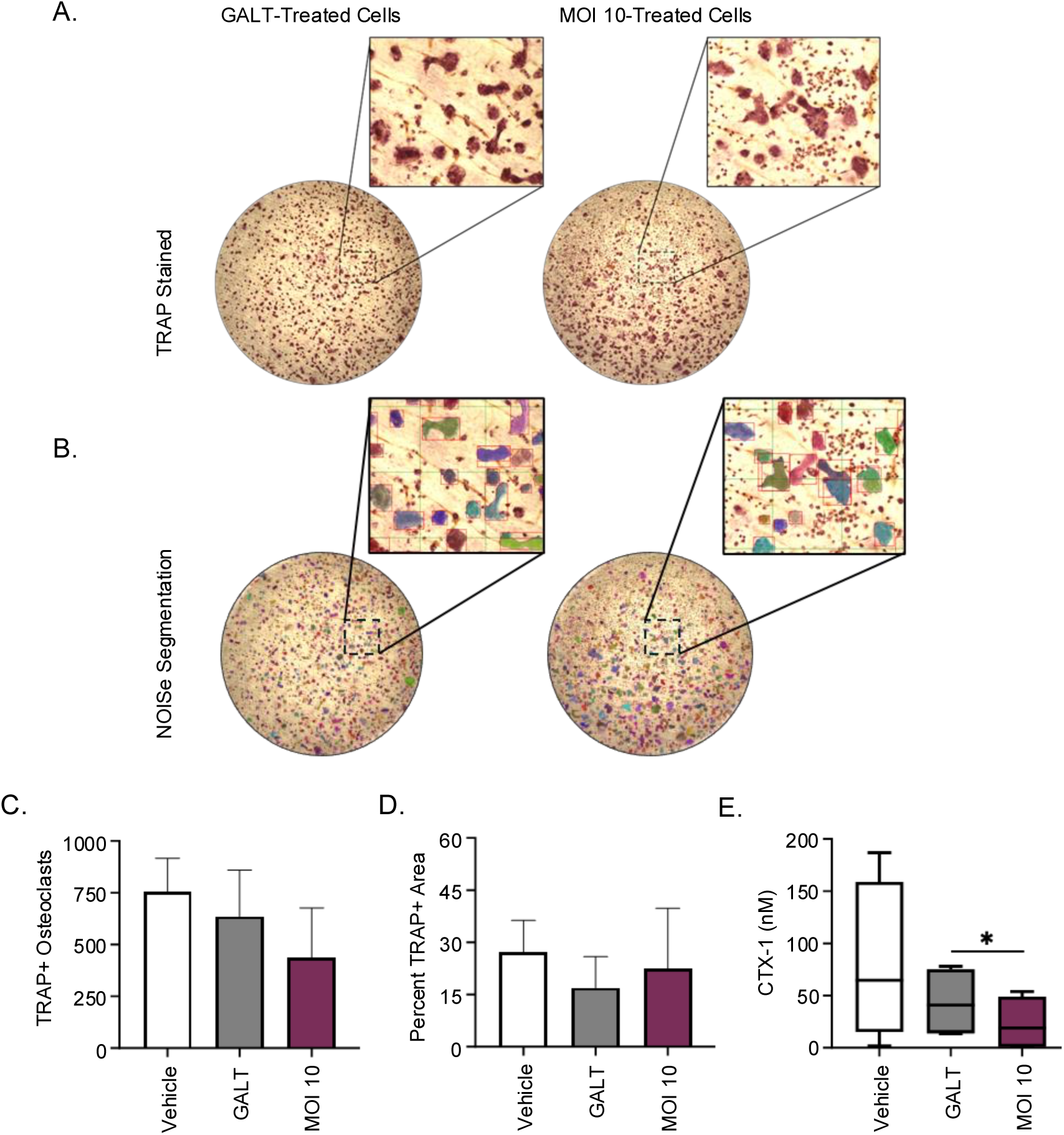
SBD111-treated media reduces the resorptive activity of osteoclasts derived from human PBMCs. Osteoclasts derived from Human PBMCs were grown on bone chips in osteoclastogenic media for 14 days with media collected at day 12 and cells were fixed and stained for TRAP at day 14. Media treatments included GALT model media (Vehicle), media control-treated GALT-conditioned media (GALT), or SBD111-treated GALT-conditioned media (MOI 10). Following fixation cells were stained for TRAP and quantified using NOISe. **(A)** Representative images of TRAP+ osteoclasts grown on bone chips illustrating effects of treated medias. **(B)** Example of NOISe quantification and segmentation abilities. **(C)** NOISe-based quantification of osteoclast number and **(D)** TRAP+ area. **(E)** CTX-1 levels (a measure of resorbed bone) were measured by ELISA in media collected from the osteoclast culture 12 days post isolation. Bars indicate mean and SEM based on n=3 for count and area measures **(C, D)**. Boxplots represent median and interquartile range, with whiskers representing max and min values, of n=4 independent experiments for bone resorption **(E)**. Vehicle condition is shown to display the normal range of human PBMC-derived osteoclast resorption and differentiation measures. Significance between conditions was determined by one-way ANOVA with Tukey’s HSD. (* = *p*<0.05).

Finally, to examine whether SBD111-GALT interactions result in factors that can inhibit bone resorption, Vehicle, GALT, and SBD111-treated GALT-conditioned (MOI 10) supernatants were added to cultures of primary human osteoclasts isolated from postmenopausal female donors and grown on bone slices for 14 days. The number of TRAP+ osteoclasts and the total TRAP+ area on each bone slice (**Fig. 4A, B, C, and D**) was not significantly different among the conditions. However, the MOI 10 supernatant significantly reduced media CTX-1 levels compared to GALT supernatant alone **(Fig. 4E),** indicating MOI 10-treated GALT media suppresses bone resorption without affecting osteoclast viability or differentiation.

## 4. Discussion

Osteopenia and osteoporosis commonly affect postmenopausal women and lead to increased fracture risk (Greendale et al., 2012; Reid & McClung, 2024). Options to mitigate bone loss are limited and, while pharmaceutical treatments exist, severe side effects limit their adoption until significant bone loss has occurred (Khan et al., 2023; MacLennan et al., 2004; Migliorati et al., 2010). To address this need, SBD111, a synbiotic medical food for the dietary management of postmenopausal bone loss was developed. In a recent double-blind, placebo-controlled clinical trial that enrolled women within 6 years of menopause, oral administration of SBD111 for one year reduced bone loss in participants with osteopenia, elevated BMI, or elevated body fat (Schott et al., 2025). Given that, SBD111 administration also corresponded with reduced serum CTX-1 in participants with BMI Δ30 and the association between elevated BMI/body fat, gut barrier dysfunction, and systemic inflammation, these findings suggest that SBD111 exerts benefits on bone health by reducing localized inflammation and bone turnover. Here, we endeavored to determine potential mechanisms through which SBD111 impacts systemic bone health.

SBD111 was shown to reduce severe GI symptoms in a human efficacy trial, indicating its direct action in the gut. As such, we hypothesized that this synbiotic medical food influences bone turnover through modulation of the intestinal barrier, a mechanism through which gastrointestinal microbes elicit broad systemic effects on immune function and health (Di Vincenzo et al., 2023a; Escalante et al., 2024). To test this, we examined its effect on intestinal epithelial monolayers and observed that SBD111 administration improved intestinal barrier functionality without inducing inflammatory chemokine secretion (IL-8 and CXCL1), independent of concentration. These phenotypes are consistent with previous reports that oral administration of beneficial microbes can reduce intestinal permeability, thereby limiting translocation of GI-resident microbes into surrounding tissues. This effect decreases local immune cell activation/inflammation and can reduce bone loss (Di Vincenzo et al., 2023a; Escalante et al., 2024; Thoo et al., 2019). Given that osteopenia and high BMI/visceral body fat are known to increase intestinal permeability and bacterial translocation, this phenotype is consistent with our clinical findings that SBD111 is effective at reducing bone loss in those populations (Ferbebouh et al., 2021; Festa et al., 2001; Kredel & Siegmund, 2014; Teixeira et al., 2012; Zhang et al., 2022). These conclusions could be strengthened by examining SBD111’s effects on damaged epithelial membranes to model the leaky gut, common to high BMI/body fat individuals. Furthermore future in vivo studies could confirm this mechanism through quantification of serum LPS or zonulin, as markers of intestinal permeability (Schoultz & Keita, 2020).

The menopause transition, obesity, and aging are all associated with increased inflammatory responses that can exacerbate bone loss (Festa et al., 2001; Kredel & Siegmund, 2014; Shieh et al., 2020; Thevaranjan et al., 2017). It has been proposed that these inflammatory responses are due, in part, to local inflammation and immune activation within the gut (Di Vincenzo et al., 2023a; Thoo et al., 2019; Zaiss et al., 2019). Given that SBD111 was most effective at reducing bone loss in a high BMI/body fat population, which are associated with systemic inflammation, we hypothesized that SBD111 administration may modulate the inflammatory responses of local immune cell populations within the gut. To test this, we examined PBMC responses to SBD111 with or without the addition of an inflammatory challenge (LPS); to mimic the inflammatory conditions associated with osteopenia and high BMI/body fat. In the absence of LPS challenge, low SBD111 concentrations induced the production of cytokines that drive T-cell differentiation/maintenance and inflammation (IL-23 and IFN-γ) and immune cell migration (CXCL1), but production of these cytokines by PBMCs decreased with higher SBD111 concentrations (Cui et al., 2025; Muranski & Restifo, 2013; Pollard et al., 2013). In the context of an inflammatory challenge, a similar concentration-dependent response was observed for IFN-γ in females, and similar but nonsignificant trends for IFN-g in males and IL-23 in females. IL-6, CXCL1, and IL-8 exhibited strong, SBD111 concentration-dependent reductions in secretion by PBMCs following inflammatory challenge. The affected cytokines are inflammatory mediators that can drive inflammation and tissue damage, and importantly, these cytokines have been shown to directly (i.e.: IL-23, IL-6, and IL-8) and indirectly (i.e.: CXCL1 and IFN-γ) enhance osteoclastogenesis (Cai et al., 2024; Capucetti et al., 2020; Gao et al., 2006; Rizo-Téllez & Filep, 2024; Umur et al., 2024). Collectively, these observations suggest that SBD111 administration reduces cytokine secretion by immune cells. However SBD111 has not been observed to alter serum cytokine responses in preclinical or human efficacy studies, suggesting that different local effects, such as the polarization and migration of immune cells, may drive SBD111s beneficial effects. It is well known that activated GI immune cells migrate throughout the body, eliciting distal effects (Di Vincenzo et al., 2023b; Galván-Peña et al., 2024). Consistently, SBD111 reduced cytokines that are critical for the polarization/maintenance of inflammatory T cells (IL-6, IFN-γ, and IL-23, non-significantly) and immune cell migration (IL-8 and CXCL1) under inflammatory challenge. As, T cell polarization is known to affect osteoclast function, this intriguing possibility warrants further investigation in preclinical studies examining immune cell activation states within GI tissues and in circulating immune cell populations (Gao et al., 2006; Sato et al., 2006).

Additionally, we noted that there was donor variation in PBMC responses to SBD111. One major source of donor variation was sex. SBD111 administration, in the context of inflammatory challenge, non-significantly decreased IL-23 and IFN-γ secretion in males, while inducing similar CXCL1, IL-8, and IL-6 responses to the females. These results suggested that males may see similar if muted benefits to SBD111; a topic that could be explored in future clinical studies. As donor variation was also seen within sexes, it would also be beneficial to explore which additional subpopulations would most benefit from SBD111. To this end, future experiments would benefit from sourcing PBMCs from donors with different immunological backgrounds (e.g., those with autoimmune/inflammatory disorders) to determine whether other inflamed populations would benefit from SBD111 administration, similar to the high BMI/body fat population examined clinically.

SBD111 has been shown to reduce serum CTX-1 levels, a marker of bone degradation, in humans with BMI ≥30, indicating that it likely inhibits osteoclast development or functioning, a process that is influenced by microbe-, epithelium-, and immune cell-derived molecules (Cai et al., 2024; Umur et al., 2024; Zaiss et al., 2019). To determine whether SBD111s interactions in the gut, and resulting products, underlie its bone sparing effects, PBMCs were cultured in the basolateral chamber of an in vitro gut model with apical administration of SBD111 (depicted in **Fig. 3A**). This is a complex multi-cellular model that contains many, but not all, of the pertinent cell types found in immunological tissues of the gut, resulting in a more physiologically relevant model capable of producing more complex responses. The SBD111-treated PBMC-conditioned media (basolateral supernatant representing the lamina propria) produced from this model was then incubated with murine RAW264.7 or human osteoclast precursor cells in the presence of osteoclast differentiating factors. Using this system, we determined that SBD111-treated basolateral supernatants reduced TRAP activity in RAW264.7 osteoclast precursor cells, indicating decreased osteoclast differentiation and/or function. Additionally, we showed that SBD111-derived factors directly inhibited TRAP activity by incubating RAW264.7 cells with SBD111-conditioned media without human cells. Similarly, we found that SBD111-conditioned PBMC supernatant administration also reduced a marker of bone resorption (CTX-1) in a more physiologically relevant in vitro model of human osteoclastogenesis and bone turnover. However, in this human primary cell model, the total number of osteoclasts (quantity of TRAP+ cells) was not affected, suggesting that SBD111 reduces osteoclast activity but not osteoclastogenesis. These observations are consistent with clinical data demonstrating that dietary intervention with SBD111 decreased serum CTX-1 in women with elevated BMI (Schott et al., 2025). It is important to note that, while this mode corroborates what is seen clinically, it does not necessarily reflect the local concentration of these products within bone which could affect the degree of inhibition seen in vivo.

As summarized in **the graphical abstract**, these data describe multifactorial mechanisms through which SBD111 medical food may confer a clinical benefit for the management of postmenopausal bone loss. We show that SBD111 improves intestinal epithelial barrier integrity and reduces inflammatory responses by immune cells in vitro, supporting our previous findings that SBD111 supplementation reduces severe GI symptoms and inflammatory signaling within bone (Lawenius et al., 2022; Schott et al., 2025). Notably, SBD111 exposure induces a concentration-dependent anti-osteoclastogenic environment in an in vitro gut model, supporting our clinical findings that dietary intervention with SBD111 reduces bone degradation markers in BMI ≥30 women (Schott et al., 2025). Together, these results provide a mechanistic foundation for further clinical investigation of the medical food, SBD111, in populations at risk for inflammation-associated bone loss.

## 5. Conclusions

SBD111, a synbiotic medical food composed of probiotic strains from fruits and vegetables as well as prebiotic fibers, has been shown to slow bone loss in women with BMI ≥30, with body fat ≥40%, or with osteopenia. Here, we rigorously tested SBD111 to elucidate potential mechanisms underlying its benefits for bone health. We have determined that SBD111 may function through multiple mechanisms, wherein it can reduce osteoclast activity (bone degradation) directly, or indirectly through reductions intestinal permeability and local inflammation.

## Acknowledgements

The authors would like to thank Clifford Rosen for advice and introducing collaborating authors, the MaineHealth Biobank for providing human whole blood for primary osteoclast culture, and Michael Wan for assistance with NOISe.

## Author Contributions

Conceptualization: RSG, DDM, AEB, and KJM; Investigation: RSG, TR, DDM, and CM; Formal analysis: RSG, TR, DDM, and RN; Writing-original draft: RSG, TR, DDM, AEB, and KJM; Writing-review and editing: RSG, TR, DDM, CM, RN, EMS, MRC, AEB, KJM, and GVT; Funding acquisition: KJM and GVT. GVT has primary responsibility for final content. All authors read and approved the final manuscript.

## Data Availability Statement

The data described in the manuscript will be made available upon request.

## IRB Statements

The study was conducted according to the guidelines of the Declaration of Helsinki and approved by the Institutional Review Board of MaineHealth under protocol 1689738-1 (approved 12/29/2020) and 958914 (approved 05/31/2005). Human peripheral blood mononuclear cells (PBMCs) were purchased from Charles River Laboratories and collected under their IRB-approved protocol with informed consent for commercial research use.

## Informed Consent Statements

Informed consent was obtained from all participants involved in the study.

## Abbreviations

αMEM (alpha Minimal Essential Medium); ΔTEER (Change in Trans-epithelial Electrical Resistance); anti-anti (antibiotic-antimycotic); ATCC (American Type Culture Collection); BMD: Bone mineral density; CFU (Colony forming units); CTX-1 (collagen cross-linked telopeptide); CXCL (C-X-C motif ligand); DMEM (Dulbecco’s Modified Eagle Medium); ELISA (Enzyme-Linked Immunosorbent Assay); FBS (Fetal Bovine Serum); GALT (gut-associated lymphoid tissue); GI (Gastrointestinal); IFN-γ (Interferon gamma); IL (Interleukin); IP-10 (IFN-γ-induced protein 10); M-CSF (Macrophage colony stimulating factor); MEM (Minimal Essential Medium); MOI (Multiplicity of interaction); OPG (Osteoprotegerin); PBMC (Peripheral blood mononuclear cells); PBS (Phosphate Buffered Saline); RANKL (Receptor activator of nuclear factor-κB Ligand); RPMI-1640 (Roswell park Memorial Institute medium); TEER (Trans-epithelial Electrical Resistance); TNF-α (Tumor Necrosis factor alpha); TRAP (Tartrate-resistant acid phosphatase); TSB (Tryptic Soy Broth)

## References

1. Boyle, W. J., Simonet, W. S., & Lacey, D. L. (2003). Osteoclast differentiation and activation. Nature 2003 *423*:6937, *423*(6937), 337–342. 10.1038/nature01658

2. Cai, L., Lv, Y., Yan, Q., & Guo, W. (2024). Cytokines: The links between bone and the immune system. Injury, 55(2), 111203. 10.1016/J.INJURY.2023.111203

3. Capucetti, A., Albano, F., & Bonecchi, R. (2020). Multiple Roles for Chemokines in Neutrophil Biology. Frontiers in Immunology, 11, 533351. 10.3389/FIMMU.2020.01259/PDF

4. Cui, X., Liu, W., Jiang, H., Zhao, Q., Hu, Y., Tang, X., Liu, X., Dai, H., Rui, H., & Liu, B. (2025). IL-12 family cytokines and autoimmune diseases: A potential therapeutic target? Journal of Translational Autoimmunity, 10, 100263. 10.1016/J.JTAUTO.2024.100263

5. Da Silva, C., Wagner, C., Bonnardel, J., Gorvel, J. P., & Lelouard, H. (2017). The Peyer’s patch mononuclear phagocyte system at steady state and during infection. In Frontiers in Immunology (Vol. 8, Issue OCT). Frontiers Media S.A. 10.3389/fimmu.2017.01254

6. Di Vincenzo, F., Del Gaudio, A., Petito, V., Lopetuso, L. R., & Scaldaferri, F. (2023a). Gut microbiota, intestinal permeability, and systemic inflammation: a narrative review. Internal and Emergency Medicine, 19(2), 275. 10.1007/S11739-023-03374-W

7. Di Vincenzo, F., Del Gaudio, A., Petito, V., Lopetuso, L. R., & Scaldaferri, F. (2023b). Gut microbiota, intestinal permeability, and systemic inflammation: a narrative review. Internal and Emergency Medicine, 19(2), 275. 10.1007/S11739-023-<otherinfo>03374-W</otherinfo>

8. Dimitrov, Z., Gotova, I., & Chorbadjiyska, E. (2014). In vitro characterization of the adhesive factors of selected probiotics to Caco-2 epithelium cell line. *Biotechnology*, Biotechnological Equipment, 28, 1079–1083. 10.1080/13102818.2014.969948

9. Easson, D. D., Murphy, V. A., Ballok, A. E., Soto-Giron, M. J., Schott, E. M., Rodricks, J., & Toledo, G. V. (2022). Food safety assessment and toxicity study of the synbiotic consortium SBD111. Food and Chemical Toxicology, 168. 10.1016/j.fct.2022.113329

10. Escalante, J., Artaiz, O., Diwakarla, S., & McQuade, R. M. (2024). Leaky gut in systemic inflammation: exploring the link between gastrointestinal disorders and age-related diseases. GeroScience 2024 *47*:1, *47*(1), 1–22. 10.1007/S11357-024-01451-2

11. Farache, J., Koren, I., Milo, I., Gurevich, I., Kim, K. W., Zigmond, E., Furtado, G. C., Lira, S. A., & Shakhar, G. (2013). Luminal Bacteria Recruit CD103+ Dendritic Cells into the Intestinal Epithelium to Sample Bacterial Antigens for Presentation. Immunity, 38(3), 581–595. 10.1016/j.immuni.2013.01.009

12. Ferbebouh, M., Vallières, F., Benderdour, M., & Fernandes, J. (2021). The pathophysiology of immunoporosis: innovative therapeutic targets. In Inflammation Research (Vol. 70, Issue 8, pp. 859–875). Springer Science and Business Media Deutschland GmbH. 10.1007/s00011-021-01484-9

13. Ferraretto, A., Bottani, M., De Luca, P., Cornaghi, L., Arnaboldi, F., Maggioni, M., Fiorilli, A., & Donetti, E. (2018). Morphofunctional properties of a differentiated Caco2/HT-29 coculture as an in vitro model of human intestinal epithelium. Bioscience Reports, 38(2). 10.1042/BSR20171497

14. Festa, A., D’Agostino, R., Williams, K., Karter, A. J., Mayer-Davis, E. J., Tracy, R. P., & Haffner, S. M. (2001). The relation of body fat mass and distribution to markers of chronic inflammation. International Journal of Obesity, 25, 1407–1415. 10.1038/sj.ijo.0801792

15. Furusawa, Y., Obata, Y., Fukuda, S., Endo, T. A., Nakato, G., Takahashi, D., Nakanishi, Y., Uetake, C., Kato, K., Kato, T., Takahashi, M., Fukuda, N. N., Murakami, S., Miyauchi, E., Hino, S., Atarashi, K., Onawa, S., Fujimura, Y., Lockett, T., … Ohno, H. (2013). Commensal microbe-derived butyrate induces the differentiation of colonic regulatory T cells. Nature 2013 504:7480, *504*(7480), 446–450. 10.1038/nature12721

16. Galván-Peña, S., Zhu, Y., Hanna, B. S., Mathis, D., & Benoist, C. (2024). A dynamic atlas of immunocyte migration from the gut. Science Immunology, 9(91). 10.1126/sciimmunol.adi0672

17. Gao, Y., Grassi, F., Ryan, M. R., Terauchi, M., Page, K., Yang, X., Weitzmann, M. N., & Pacifici, R. (2006). IFN-γ stimulates osteoclast formation and bone loss in vivo via antigen-driven T cell activation. Journal of Clinical Investigation, 117(1), 122. 10.1172/JCI30074

18. Greendale, G. A., Sowers, M., Han, W., Huang, M. H., Finkelstein, J. S., Crandall, C. J., Lee, J. S., & Karlamangla, A. S. (2012). Bone Mineral Density Loss in Relation to the Final Menstrual Period in a Multi-ethic Cohort: Results from the Study of Women’s Health Across the Nation (SWAN). Journal of Bone and Mineral Research : The OTicial Journal of the American Society for Bone and Mineral Research, 27(1), 111. 10.1002/JBMR.534

19. Hosmer, J., McEwan, A. G., & Kappler, U. (2024). Bacterial acetate metabolism and its influence on human epithelia. Emerging Topics in Life Sciences, 8(1), 1–13. 10.1042/ETLS20220092

20. Hua, M. C., Lin, T. Y., Lai, M. W., Kong, M. S., Chang, H. J., & Chen, C. C. (2010). Probiotic Bio-Three induces Th1 and anti-inflammatory effects in PBMC and dendritic cells. World Journal of Gastroenterology : WJG, 16(28), 3529. 10.3748/WJG.V16.I28.3529

21. Janský, L., Reymanová, P., & Kopecký, J. (2003). Dynamics of Cytokine Production in Human Peripheral Blood Mononuclear Cells Stimulated by LPS or Infected by Borrelia. Physiol. Res, 52, 593–598. http://www.biomed.cas.cz/physiolres

22. Khan, S. J., Kapoor, E., Faubion, S. S., & Kling, J. M. (2023). Vasomotor Symptoms During Menopause: A Practical Guide on Current Treatments and Future Perspectives. International Journal of Women’s Health, 15, 273–287. 10.2147/IJWH.S365808

23. Kleiveland, C., & Kleiveland, C. (2015). Peripheral Blood Mononuclear Cells. The Impact of Food Bioactives on Health: In Vitro and Ex Vivo Models, 161–167. 10.1007/978-3-319-16104-4_15

24. Kleiveland, C. R. (2015). Co-culture Caco-2/Immune Cells. The Impact of Food Bioactives on Health: In Vitro and Ex Vivo Models, 197–205. 10.1007/978-3-319-16104-4_18

25. Korsten, S. G. P. J., Vromans, H., Garssen, J., & Willemsen, L. E. M. (2023). Butyrate Protects Barrier Integrity and Suppresses Immune Activation in a Caco-2/PBMC Co-Culture Model While HDAC Inhibition Mimics Butyrate in Restoring Cytokine-Induced Barrier Disruption. Nutrients, 15(12). 10.3390/nu15122760

26. Kredel, L. I., & Siegmund, B. (2014). Adipose-tissue and intestinal inflammation - visceral obesity and creeping fat. Frontiers in Immunology, 5(SEP), 112587. 10.3389/FIMMU.2014.00462/PDF

27. Kumar, S., Manne, R., Martin, B., Roy, T., Neilson, R., Peters, R., Chillara, M., Lary, C. W., Motyl, K. J., & Wan, M. (2024). NOISe: Nuclei-Aware Osteoclast Instance Segmentation for Mouse-to-Human Domain Transfer. Conf Comput Vis Pattern Recognit Workshops. 10.1109/cvprw63382.2024.00686

28. Lawenius, L., Gustafsson, K. L., Wu, J., Nilsson, K. H., Movérare-Skrtic, S., Schott, E. M., Soto-Girón, M. J., Toledo, G. V., Sjögren, K., & Ohlsson, C. (2022). Development of a synbiotic that protects against ovariectomy-induced trabecular bone loss. American Journal of Physiology - Endocrinology and Metabolism, 322(4), E344–E354. 10.1152/AJPENDO.00366.2021

29. Li, J. Y., Chassaing, B., Tyagi, A. M., Vaccaro, C., Luo, T., Adams, J., Darby, T. M., Weitzmann, M. N., Mulle, J. G., Gewirtz, A. T., Jones, R. M., & Pacifici, R. (2016). Sex steroid deficiency–associated bone loss is microbiota dependent and prevented by probiotics. The Journal of Clinical Investigation, 126(6), 2049. 10.1172/JCI86062

30. Li, S., Liu, G., & Hu, S. (2024). Osteoporosis: interferon-gamma-mediated bone remodeling in osteoimmunology. Frontiers in Immunology, 15, 1396122. 10.3389/FIMMU.2024.1396122/BIBTEX

31. MacLennan, A. H., Broadbent, J. L., Lester, S., & Moore, V. (2004). Oral oestrogen and combined oestrogen/progestogen therapy versus placebo for hot flushes. In Cochrane Database of Systematic Reviews (Vol. 2009). John Wiley and Sons Ltd. 10.1002/14651858.CD002978.pub2

32. Mazziotta, C., Tognon, M., Martini, F., Torreggiani, E., & Rotondo, J. C. (2023). Probiotics Mechanism of Action on Immune Cells and Beneficial Effects on Human Health. In Cells (Vol. 12, Issue 1). MDPI. 10.3390/cells12010184

33. Migliorati, C. A., Woo, S. Bin, Hewson, I., Barasch, A., Elting, L. S., Spijkervet, F. K. L., & Brennan, M. T. (2010). A systematic review of bisphosphonate osteonecrosis (BON) in cancer. Supportive Care in Cancer, 18(8), 1099–1106. 10.1007/S00520-010-0882-1/METRICS

34. Moles, L., & Otaegui, D. (2020). The impact of diet on microbiota evolution and human health. Is diet an adequate tool for microbiota modulation? In Nutrients (Vol. 12). MDPI AG. 10.3390/nu12061654

35. Møller, A. M. J., Delaissé, J. M., Olesen, J. B., Madsen, J. S., Canto, L. M., Bechmann, T., Rogatto, S. R., & Søe, K. (2020). Aging and menopause reprogram osteoclast precursors for aggressive bone resorption. Bone Research 2020 8:1, *8*(1), 1–11. 10.1038/s41413-020-0102-7

36. Muranski, P., & Restifo, N. P. (2013). Essentials of Th17 cell commitment and plasticity. Blood, 121(13), 2402. 10.1182/BLOOD-2012-09-378653

37. Ngkelo, A., Meja, K., Yeadon, M., Adcock, I., & Kirkham, P. A. (2012). LPS induced inflammatory responses in human peripheral blood mononuclear cells is mediated through NOX4 and Giα dependent PI-3kinase signalling. *Journal of Inflammation (London*, England), 9, 1. 10.1186/1476-9255-9-1

38. Pal, S., Morgan, X., Dar, H. Y., Gacasan, C. A., Patil, S., Stoica, A., Hu, Y. J., Weitzmann, M. N., Jones, R. M., & Pacifici, R. (2024). Gender-affirming hormone therapy preserves skeletal maturation in young mice via the gut microbiome. Journal of Clinical Investigation, 134(10). 10.1172/JCI175410

39. Pandey, K. R., Naik, S. R., & Vakil, B. V. (2015). Probiotics, prebiotics and synbiotics- a review. In Journal of Food Science and Technology (Vol. 52, Issue 12, pp. 7577–7587). Springer India. 10.1007/s13197-015-1921-1

40. Pollard, K. M., Cauvi, D. M., Toomey, C. B., Morris, K. V., & Kono, D. H. (2013). Interferon-γ and Systemic Autoimmunity. Discovery Medicine, 16(87), 123. https://pmc.ncbi.nlm.nih.gov/articles/PMC3934799/

41. Quach, D., Parameswaran, N., McCabe, L., & Britton, R. A. (2019). Characterizing how probiotic Lactobacillus reuteri 6475 and lactobacillic acid mediate suppression of osteoclast differentiation. Bone Reports, 11, 100227. 10.1016/J.BONR.2019.100227

42. Rahman, M., Kukita, A., Kukita, T., Shobuike, T., Nakamura, T., & Kohashi, O. (2003). Two histone deacetylase inhibitors, trichostatin A and sodium butyrate, suppress differentiation into osteoclasts but not into macrophages. Blood, 101(9), 3451–3459. 10.1182/BLOOD-2002-08-2622

43. Reid, I. R., & McClung, M. R. (2024). Osteopenia: a key target for fracture prevention. The Lancet Diabetes and Endocrinology, 12(11), 856–864. 10.1016/S2213-8587(24)00225-0/ASSET/AD1E2EF1-65CE-45CB-96AC-99A45ED2044E/MAIN.ASSETS/GR4.SML

44. Ren, D., Wang, D., Liu, H., Shen, M., & Yu, H. (2019). Two strains of probiotic Lactobacillus enhance immune response and promote naive T cell polarization to Th1. Food and Agricultural Immunology, 30(1), 281–295. 10.1080/09540105.2019.1579785

45. Rizo-Téllez, S. A., & Filep, J. G. (2024). Beyond host defense and tissue injury: the emerging role of neutrophils in tissue repair. In American Journal of Physiology - Cell Physiology (Vol. 326, pp. C661–C683). American Physiological Society. 10.1152/ajpcell.00652.2023

46. Sahni, S., Schott, E. M., Carroll, D., Soto-Giron, M. J., Corbett, S., V Toledo, G., & P Kiel, D. (2023). Randomized clinical trial to test the safety and tolerability of SBD111, an optimized synbiotic medical food combination designed for the dietary management of the metabolic processes underlying osteopenia and osteoporosis. Journal of Microbiology & Experimentation, 11(1), 1–11. 10.15406/jmen.2023.11.00379

47. Sato, K., Suematsu, A., Okamoto, K., Yamaguchi, A., Morishita, Y., Kadono, Y., Tanaka, S., Kodama, T., Akira, S., Iwakura, Y., Cua, D. J., & Takayanagi, H. (2006). Th17 functions as an osteoclastogenic helper T cell subset that links T cell activation and bone destruction. The Journal of Experimental Medicine, 203(12), 2673. 10.1084/JEM.20061775

48. Schott, E. M., Charbonneau, M., Kiel, D. P., Bukata, S., Zuscik, M. J., Rosen, C., Ballok, A., Toledo, G. V., Steels, E., Huntress, H., Rao, A., Travison, T. G., Soto-Giron, M. J., Wolff, I., Easson, D. D., Engelke, K., & Vitetta, L. (2025). A randomized, double-blind, placebo-controlled clinical study to evaluate the efficacy of the synbiotic medical food, SBD111, for the clinical dietary management of bone loss in menopausal women. Osteoporosis International, In Press. 10.1101/2025.05.20.25325893

49. Schoultz, I., & Keita, Å. V. (2020). The Intestinal Barrier and Current Techniques for the Assessment of Gut Permeability. In Cells (Vol. 9, Issue 8). NLM (Medline). 10.3390/cells9081909

50. Scotland, R. S., Stables, M. J., Madalli, S., Watson, P., & Gilroy, D. W. (2011). Sex differences in resident immune cell phenotype underlie more efficient acute inflammatory responses in female mice. Blood, 118(22), 5918–5927. 10.1182/blood-2011-03-340281

51. Shieh, A., Epeldegui, M., Karlamangla, A. S., & Greendale, G. A. (2020). Gut permeability, inflammation, and bone density across the menopause transition. JCI Insight, 5(2), e134092. 10.1172/JCI.INSIGHT.134092

52. Šromová, V., Sobola, D., & Kaspar, P. (2023). A Brief Review of Bone Cell Function and Importance. Cells, 12(21), 2576. 10.3390/CELLS12212576

53. Takegahara, N., Kim, H., & Choi, Y. (2024). Unraveling the intricacies of osteoclast differentiation and maturation: insight into novel therapeutic strategies for bone-destructive diseases. Experimental & Molecular Medicine 2024 56:2, 56(2), 264–272. 10.1038/s12276-024-01157-7

54. Teixeira, T. F. S., Collado, M. C., Ferreira, C. L. L. F., Bressan, J., & Peluzio, M. do C. G. (2012). Potential mechanisms for the emerging link between obesity and increased intestinal permeability. Nutrition Research, 32(9), 637–647. 10.1016/J.NUTRES.2012.07.003

55. Thevaranjan, N., Puchta, A., Schulz, C., Naidoo, A., Szamosi, J. C., Verschoor, C. P., Loukov, D., Schenck, L. P., Jury, J., Foley, K. P., Schertzer, J. D., Larché, M. J., Davidson, D. J., Verdú, E. F., Surette, M. G., & Bowdish, D. M. E. (2017). Age-Associated Microbial Dysbiosis Promotes Intestinal Permeability, Systemic Inflammation, and Macrophage Dysfunction. Cell Host & Microbe, 21(4), 455. 10.1016/J.CHOM.2017.03.002

56. Thoo, L., Noti, M., & Krebs, P. (2019). Keep calm: the intestinal barrier at the interface of peace and war. In Cell Death and Disease (Vol. 10, Issue 11). Springer Nature. 10.1038/s41419-019-2086-z

57. Tyagi, A. M., Yu, M., Darby, T. M., Vaccaro, C., Li, J. Y., Owens, J. A., Hsu, E., Adams, J., Weitzmann, M. N., Jones, R. M., & Pacifici, R. (2018). The Microbial Metabolite Butyrate Stimulates Bone Formation via T Regulatory Cell-Mediated Regulation of WNT10B Expression. Immunity, 49(6), 1116. 10.1016/J.IMMUNI.2018.10.013

58. Umur, E., Bulut, S. B., Yiğit, P., Bayrak, E., Arkan, Y., Arslan, F., Baysoy, E., Kaleli-Can, G., & Ayan, B. (2024). Exploring the Role of Hormones and Cytokines in Osteoporosis Development. Biomedicines 2024, Vol. 12, Page 1830, 12(8), 1830. 10.3390/BIOMEDICINES12081830

59. White, A. A., Lin, A., Bickendorf, X., Cavve, B. S., Moore, J. K., Siafarikas, A., Strickland, D. H., & Leffler, J. (2022). Potential immunological effects of gender-affirming hormone therapy in transgender people – an unexplored area of research. In Therapeutic Advances in Endocrinology and Metabolism (Vol. 13). SAGE Publications Ltd. 10.1177/20420188221139612

60. Yuan, L., van der Mei, H. C., Busscher, H. J., & Peterson, B. W. (2020). Two-Stage Interpretation of Changes in TEER of Intestinal Epithelial Layers Protected by Adhering Bifidobacteria During E. coli Challenges. Frontiers in Microbiology, 11. 10.3389/fmicb.2020.599555

61. Zaiss, M. M., Jones, R. M., Schett, G., & Pacifici, R. (2019). The gut-bone axis: how bacterial metabolites bridge the distance. The Journal of Clinical Investigation, 129(8), 3018. 10.1172/JCI128521

62. Zhang, W., Gao, R., Rong, X., Zhu, S., Cui, Y., Liu, H., & Li, M. (2022). Immunoporosis: Role of immune system in the pathophysiology of different types of osteoporosis. Frontiers in Endocrinology, 13, 965258. 10.3389/FENDO.2022.965258

