## Supplementary Methods and Data for "A synbiotic medical food improves gut barrier function, reduces immune responses, and inhibits osteoclast activity in models of postmenopausal bone loss aligned with clinical outcomes"

### qRT-PCR

Total RNA was isolated from pooled GALT model epithelial cells and PBMCs using a commercially available kit (*Quick*-RNA™ Miniprep Plus Kit; CAT# R1058; Zymo Research, Irvine, CA). Isolated mRNA was examined by qRT-PCR to determine the relative gene expression of *tnfsf11* (RANKL) and *tnfrsf11b* (OPG). RNA was converted to cDNA and quantified by real-time PCR on a Biorad CFX96 Touch Real-Time PCR detection system using PrimeTime™ 1-step broad range qPCR master mix (CAT# 10011744, Integrated DNA Technologies (IDT), Coralville, IA) and commercially available primer-probe sets (IDT, Coralville, IA. *tnfsf11: Hs.PT.58.23324760; tnfrsf11b: Hs.PT.58.24746239; gapdh: HSs.PT.39a.22214836).* Primer sequences may be found in **Table 1.** Relative gene expression was determined via GAPDH normalization using the $2^{-\Delta\Delta Ct}$ method (1).

**Table 1: Primers used in this study**

| Designation | Sequence |
|---|---|
| *tnfrsf11b (OPG)* Primer 1 | CTACCAAGACACTAAGCCAGT |
| *tnfrsf11b (OPG)* Primer 2 | AAACAGTGAATCAACTCAAAAATGTG |

| | |
|---|---|
| *tnfrsf11b (OPG)* Probe | /56-<br>FAM/AACCTGAAG/ZEN/AATGCCTCCTCACACAG/3IABkFQ/ |
| *tnfsf11 (RANKL)*<br>Primer 1 | TGAGATGAGCAAAAGGCTGAG |
| *tnfsf11 (RANKL)*<br>Primer 2 | AGGAGCTGTGCAAAAGGAAT |
| *tnfsf11 (RANKL)* Probe | /5HEX/CCAGATCTA/ZEN/ACCATGAGCCATCCACC/3IABkFQ/ |
| *gapdh* Primer 1 | TGTAGTTGAGGTCAATGAAGGG |
| *gapdh* Primer 2 | ACATCGCTCAGACACCATG |
| *gapdh* Probe | /5Cy5/AAGGTCGGA/TAO/GTCAACGGATTTGGTC/3IAbRQSp/ |

1. Livak, K. J. & Schmittgen, T. D. Analysis of Relative Gene Expression Data Using Real-Time Quantitative PCR and the 2−ΔΔCT Method. *Methods* **25**, 402–408 (2001).

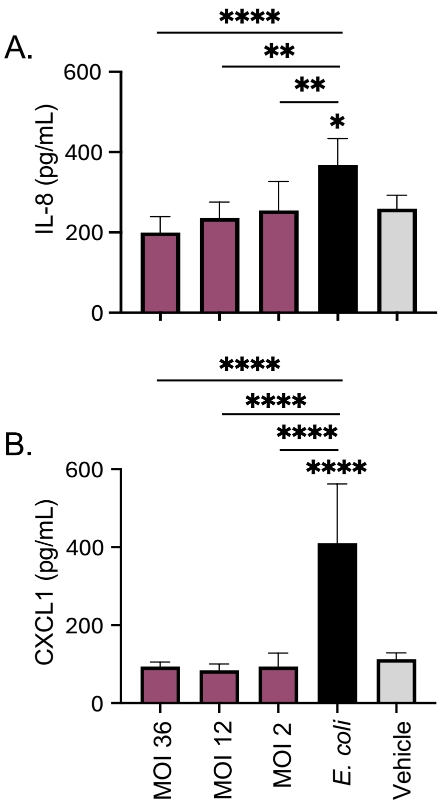


**Supplementary Figures:**

**Supplementary Figure 1. SBD111 does not induce inflammatory chemokine secretion from Caco-2 and HT29 cell monolayers.** The mature, polarized intestinal epithelial cell (Caco-2 and HT29) monolayers exposed to a media control (Vehicle), disruption control (*E. coli*), or SBD111 material at MOIs of 36, 12, or 2 for 24 h. After 24h **(A)** IL-8 and **(B)** CXCL1 secretion was analyzed by ELISA. These data correspond to the experiment presented in **Fig. 1** and are representative of eight experiments. Results are expressed as mean + SD of 12 replicates per condition. Significance between conditions was determined by one-way ANOVA with Tukey's HSD. Asterisks without comparison bars indicate significance relative to the vehicle control. (* = $p$<0.05, ** = $p$<0.005, **** = $p$< 0.0001)

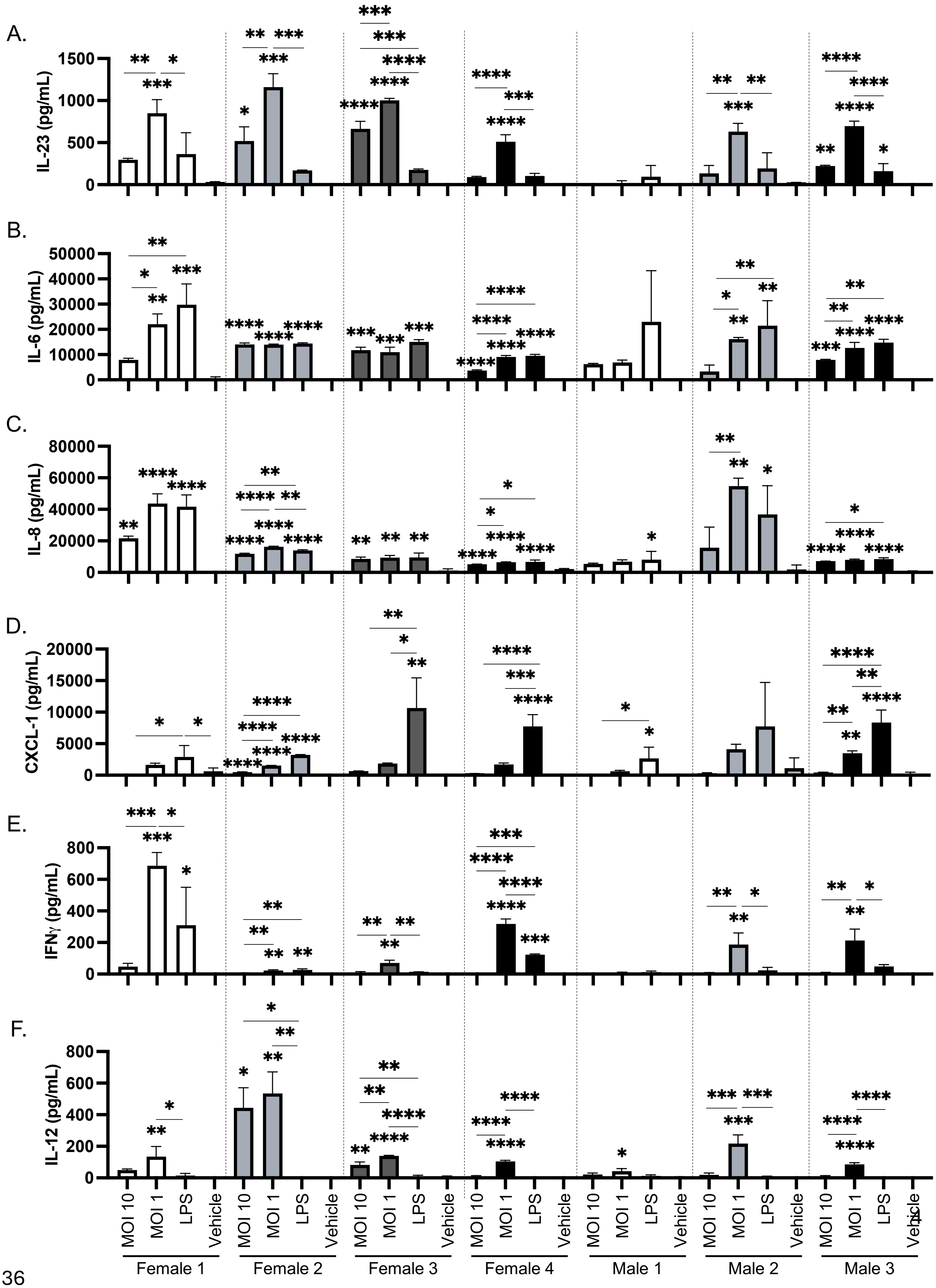
A.
IL-23 (pg/mL)
B.
IL-6 (pg/mL)
C.
IL-8 (pg/mL)
D.
CXCL-1 (pg/mL)
E.
IFNγ (pg/mL)
F.
IL-12 (pg/mL)
MOI 10
MOI 1
LPS
Vehicle
Female 1
Female 2
Female 3
Female 4
Male 1
Male 2
Male 3

**Supplementary Figure 2: SBD111 induces concentration-dependent reductions in inflammatory responses across multiple donors at baseline** This figure displays the inflammation-naïve data shown in **Fig. 2A, C, E, G,** and **I** separated by donor without LPS normalization. Briefly human PBMCs from four peri- and postmenopausal females and three male donors were treated with a vehicle control (media), LPS, or SBD111 material at an MOI of 10 or 1 for 24 h. Absolute cytokine secretion was determined by ELSIA: **(A)** IL-23, **(B)** IL-6, **(C)** IL-8, **(D)** CXCL1, **(E)** IFN–$\gamma$, and **(F)** IL-12. Graphed values are indicative of the mean + SD for a single experiment that is representative of each donor's response. Significance between conditions within the samples from an individual donor was determined by one-way ANOVA with Tukey's HSD. Asterisks without comparison bars indicate significance relative to the vehicle control. (* = $p$<0.05, ** = $p$<0.005, *** = $p$<0.001, **** = $p$<0.0001).

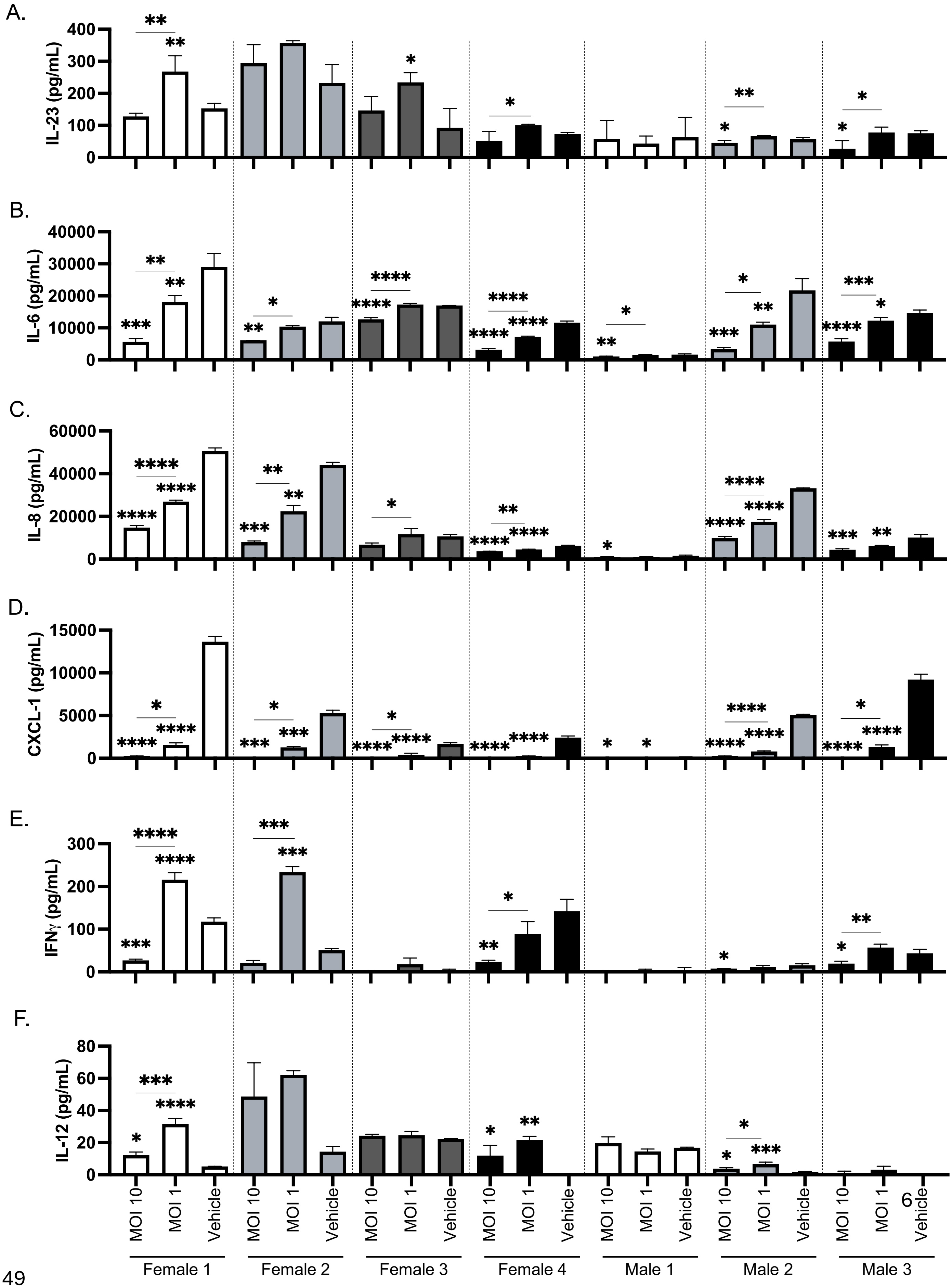
A.
IL-23 (pg/mL)
B.
IL-6 (pg/mL)
C.
IL-8 (pg/mL)
D.
CXCL-1 (pg/mL)
E.
IFNγ (pg/mL)
F.
IL-12 (pg/mL)
MOI 10
MOI 1
Vehicle
Female 1
Female 2
Female 3
Female 4
Male 1
Male 2
Male 3

**Supplementary Figure 3: SBD111 induces concentration-dependent reductions in inflammatory responses across multiple donors after an inflammatory challenge**

These data correspond to the LPS normalized data presented in **Fig. 2B, D, F, H,** and **J** without LPS normalization and separated by donor. Human PBMCs from the same four peri- and postmenopausal females and three males were challenged with 100 ng/mL of LPS for 30 min to induce inflammatory responses. After which, each sample was washed with 1 X PBS and resuspended in cell culture media and were incubated with a vehicle control (media) or SBD111 material (MOI of 10 or 1) for 24 h and absolute cytokine secretion was quantified by ELISA **(A)** IL-23, **(B)** IL-6, **(C)** IL-8, **(D)** CXCL1, **(E)** IFN–$\gamma$, and **(F)** IL-12. Each column indicates the mean + SD for an individual donor. Significance between conditions was determined within individual donors by one-way ANOVA with Tukey's HSD. Asterisks without comparison bars indicate significance relative to the vehicle control. (* = $p$<0.05, ** = $p$<0.005, *** = $p$<0.001, **** = $p$<0.0001).

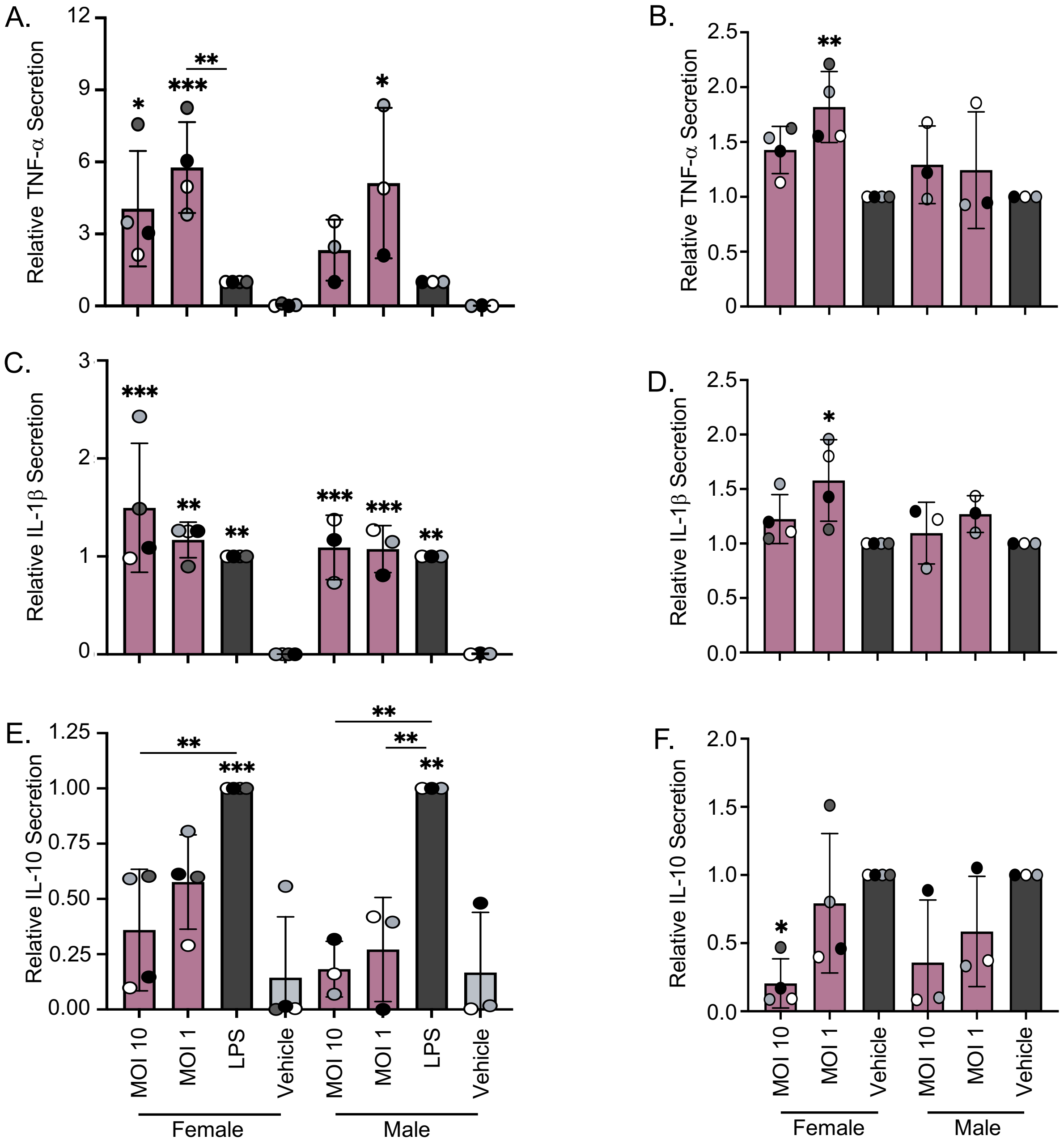


**Supplementary Figure 4: Additional cytokine responses elicited by SBD111-treated human PBMCs at baseline and after inflammatory challenge. (A, C, and E)** Human PBMCs from four peri- and postmenopausal female and three male donors were incubated with a media control (vehicle), LPS, or SBD111 material (MOI of 10 or 1) for 24 h and immune responses were determined. **(B, D, F, H, and J) Alternatively,**

PBMCs from the same donors were treated with 100 ng/mL of LPS for 30 min to induce inflammatory responses and washed with 1 X PBS. These inflamed cells were then treated with a vehicle control or SBD111 material ( MOI of 10 or 1) for 24 h. Cytokine responses were quantified via ELISA, **(A and B)** IL-23, **(C and D)** IL-6, **(E and F)** IL-8, **(G and H)** CXCL1, and **(I and J)** IFN–γ. Data was normalized to LPS (inflammation naïve conditions) or LPS-challenged, vehicle-treated controls (inflammatory challenge) to compare across donors. Columns indicate the mean + SD for all donors, differentiated by sex. Points indicate the mean response of an individual donor and are color coded by donor. Significance between conditions was determined within each sex by one-way ANOVA with Tukey's HSD. Asterisks without comparison bars indicate significance relative to the vehicle control. (* = $p<0.05$, ** = $p<0.005$, *** = $p<0.001$, **** = $p<0.0001$).

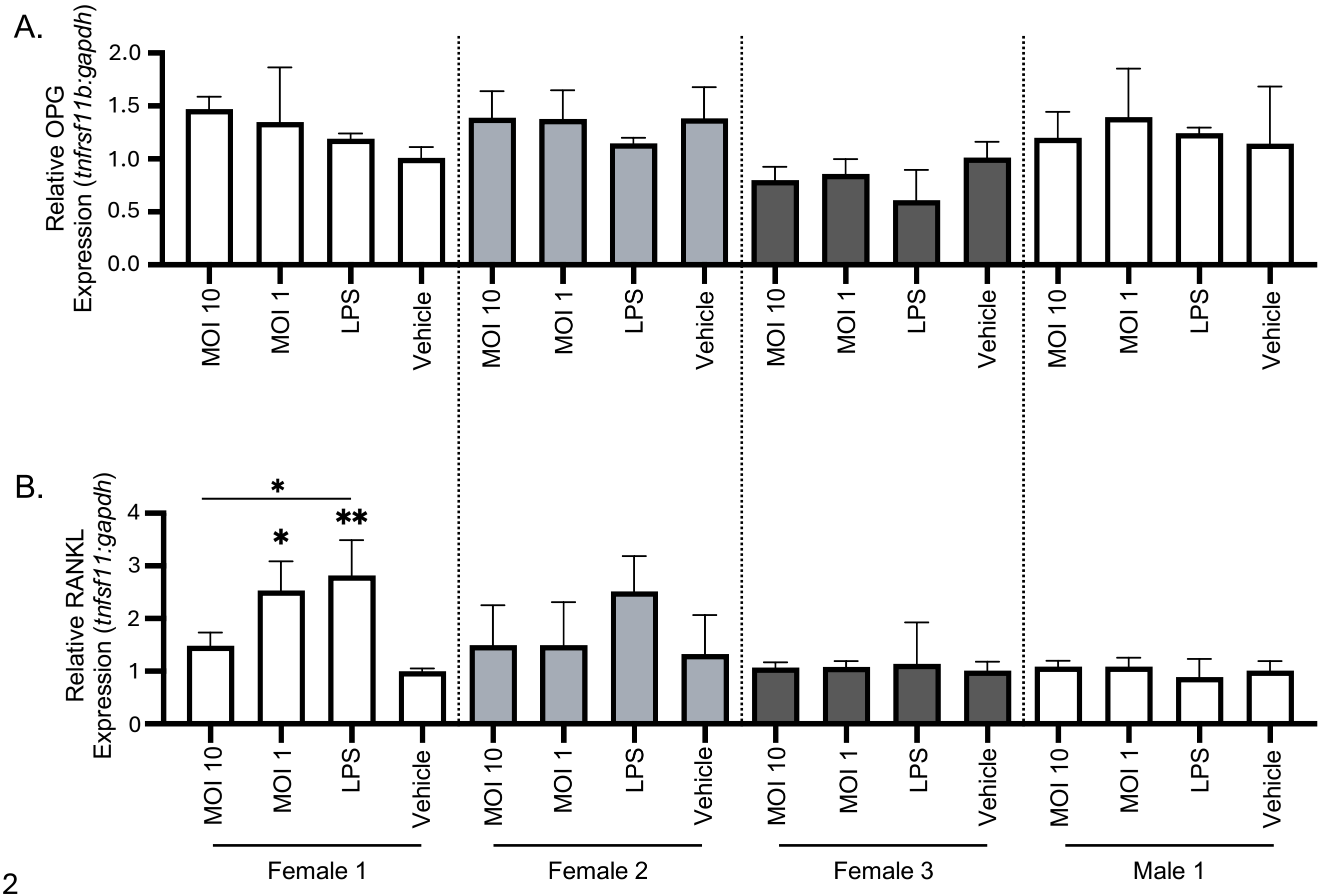


**Supplementary Figure 5: SBD111 does not alter the expression of RANKL and OPG in a GALT model.** GALT models were established, containing polarized intestinal epithelial cell (Caco-2 and HT29) monolayers on cell culture inserts with human PBMCs in the basolateral compartment. A media control, LPS (a stimulatory control), or SBD111 material at MOIs of 2, 10, or 50 were added to the apical compartment of the GALT model for 24 h. The human cells in these models were harvested and their total RNA was isolated. The relative gene expression of **(A)** osteoprotegerin (OPG; *tnfrsf11b*) and **(B)** RANKL (*tnfrsf11*) were determined by qRT-PCR. Relative gene expression was determined relative to *gapdh* by the $2^{-\Delta\Delta Ct}$ method[1]. Graphed values indicate the mean + SD for a single experiment that is representative of at least three experiments for each donor. Significance between conditions within an individual donor's samples was determined by one-way ANOVA with Tukey's HSD. Asterisks without comparison bars indicate significance relative to the vehicle control. (* = $p<0.05$, ** = $p<0.005$)
